# KAS-ATAC reveals the genome-wide single-stranded accessible chromatin landscape of the human genome

**DOI:** 10.1101/2024.05.06.591268

**Authors:** Samuel H. Kim, Georgi K. Marinov, William J. Greenleaf

**Author notes:** These authors contributed equally to this work (Samuel H. Kim, Georgi K. Marinov).

## Abstract

Gene regulation in most eukaryotes involves two fundamental physical processes – alterations in the packaging of the genome by nucleosomes, with active *cis*-regulatory elements (CREs) generally characterized by an open-chromatin configuration, and the activation of transcription. Mapping these physical properties and biochemical activities genome-wide – through profiling chromatin accessibility and active transcription – are key tools used to understand the logic and mechanisms of transcription and its regulation. However, the relationship between these two states has until now not been accessible to simultaneous measurement. To address this, we developed KAS-ATAC, a combination of the KAS-seq (Kethoxal-Assisted SsDNA sequencing and ATAC-seq (Assay for Transposase-Accessible Chromatin using sequencing) methods for mapping single-stranded DNA (and thus active transcription) and chromatin accessibility, respectively, enabling the genome-wide identification of DNA fragments that are simultaneously accessible and contain ssDNA. We use KAS-ATAC to evaluate levels of active transcription over different classes of regulatory elements in the human genome, to estimate the absolute levels of transcribed accessible DNA over CREs, to map the nucleosomal configurations associated with RNA polymerase activities, and to assess transcription factor association with transcribed DNA through transcription factor binding site (TFBS) footprinting. We observe lower levels of transcription over distal enhancers compared to promoters, surprisingly high abundance of ssDNA immediately around/within CTCF occupancy footprints, and distinct nucleosomal configurations around transcription initiation sites associated with active transcription. Remarkably, most TFs associate equally with transcribed and non-transcribed DNA but a few factors specifically do not exhibit footprints over ssDNA-containing fragments. We anticipate KAS-ATAC to continue to derive useful insights into chromatin organization and transcriptional regulation in other contexts in the future.

## Introduction

Active *cis*-regulatory elements in most eukaryotes are usually characterized by low levels of nucleosome occupancy, a unique property first appreciated more than four decades ago ^1–3^. Therefore, active CREs are found in regions of increased chromatin accessibility. The genome-wide profiling of chromatin accessibility has been a key tool for identifying active regulator elements and assessing their dynamics across time and space. Methods for carrying out this mapping use the preferential cleavage or labeling of physically accessible DNA by various enzymes. Cleavage by DNase I was the first such approach to be widely adopted, initially coupled to microarray readouts ^4,5^ and later combined with high-throughput sequencing in the form of DNase-seq ^6–8^. An alternative approach has been to use methylation and either short- or long-read readouts to map chromatin accessibility both genome-wide and at a single-molecule level ^9–11^. The most widely used assay for mapping open chromatin is ATAC-seq ^12^, which is based on the preferential insertion of the hyperactive Tn5 transposase into accessible DNA.

Apart from identifying putative regulatory elements, ATAC-seq can also be used to map nucleosome positioning in the vicinity of CREs ^13^, and both DNase-seq and ATACseq can also footprint the interactions between individual transcription factors (TFs) and DNA, as TFs protect DNA from cleavage/insertion ^14–20^.

Gene regulatory inputs in the form of transcription factor occupancy and chromatin remodeler activity are integrated into changes in transcriptional activity at promoters. Therefore, mapping active transcription has also long been a key objective (contrasted with conventional RNA-seq profiling of steady-state polyadenylated transcripts ^21–24^, the levels of which reflect several additional layers of regulation of RNA stability).

To this end, several methods for mapping nascent transcription, based on adaptations of the nuclear run-on techniques or capturing RNA polymerase molecules associated with DNA, such as NET-seq ^27^, GRO-seq ^28,29^, PRO-seq ^30^, and variations of ^31^ have been developed. However, these methods often involve very elaborate experimental protocols and thus their deployment has been relatively limited. The recently developed KAS-seq assay ^32^, by contrast, provides a straightforward method for identifying RNA polymerase-associated DNA based on the highly specific covalent labeling of unpaired single-stranded guanine residues with N_3_-kethoxal (Figure 1a); N_3_-kethoxal adducts can subsequently be subjected to a click reaction and the addition of biotin, which is then used to specifically pull down labeled DNA fragments. Most ssDNA in the genome is found in the context of the transcriptional bubbles of elongating and paused RNA polymerases, with the rest arising from active replication and secondary structures such as G-quadruplexes ^33^.

**Figure 1:**
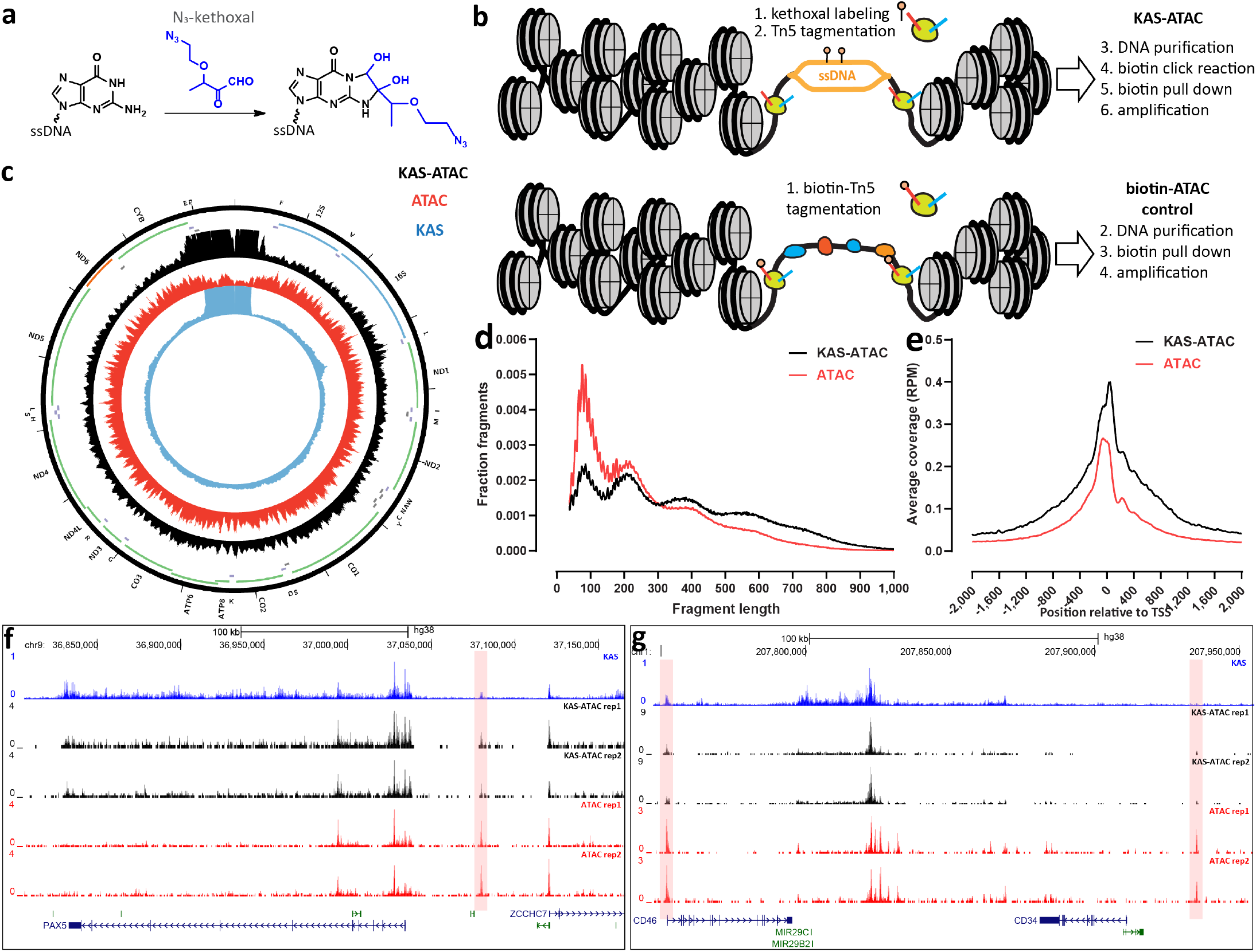
Overview of the KAS-ATAC assay. (a) N_3_-kethoxal specifically covalently labels unpaired guanine bases. (b) KAS-ATAC captures DNA fragments that are simultaneously single-stranded and accessible, by first labeling ss-DNA with N_3_-kethoxal, then carrying out an ATAC-seq reaction. After purification of DNA and a click reaction that conjugates biotin onto kethoxal-labeled DNA, accessible ssDNA is specifically pulled down using streptavidin beads and amplified. In parallel, a control “biotin-ATAC” library can be prepared using biotinylated Tn5 that then undergoes the same pull-down procedure as KAS-ATAC DNA fragments (if the goal is to quantify the relative abundances of accessible ssDNA and accessible DNA). (c) KAS-seq, ATAC-seq, and KAS-ATAC mitochondrial genome profiles in human GM12878 cells. (d) Fragment length distribution in biotin-ATAC-seq and KAS-ATAC libraries (GM12878 cells). (e) Genome-wide TSS metaprofiles for biotin-ATAC-seq and KAS-ATAC libraries (GM12878 cells). (f,g) Representative genome browser snapshots showing ATAC-seq, KAS-seq, and KAS-ATAC profiles around nuclear loci in GM12878 cells.

Unlike RNA-based methods for mapping RNA polymerase activity, KAS-seq provides the unique opportunity to directly map the relationship between chromatin accessibility and polymerase activity (and potentially other ssDNA structures), i.e. to what extent are various active CREs subject to active transcription, what nucleosomal configurations is this activity associated with, and what transcription factors may be enriched or depleted in terms of their physical associations with DNA on actively transcribed DNA fragments, as kethoxal labels the same DNA molecule that accessibility is measured on rather than a separate RNA one.

To address these questions, we developed KAS-ATAC, a method for jointly profiling ssDNA and chromatin accessibility (Figure 1b). We applied KAS-ATAC together with appropriate controls to evaluate the relative degree of transcription in different classes of regulatory elements and chromatin states in the human genome as well as to estimate the absolute abundance of transcribed and accessible/single-stranded structures. In addition, we map nucleosomal and subnucleosomal structures associated with active transcription and comprehensively measure TF footprints on polymerase-bound DNA molecules.

## Results

### KAS-ATAC maps identify genomic DNA fragments that are simultaneously accessible and contain single-stranded DNA

We developed KAS-ATAC by taking advantage of the complementary properties of the Tn5 transposase, which preferentially inserts into open chromatin, and the N_3_-kethoxal compound, which covalently attaches to unpaired G bases in ssDNA with very high specificity (Figure 1a). N_3_-kethoxal adducts can then be subjected to click chemistry and biotin addition, and DNA fragments thus labeled can be stringently purified using streptavidin pulldown. To capture DNA fragments that are simultaneously accessible and containing ssDNA, we combined these two properties into KAS-ATAC as follows (Figure 1b). First, cells are subjected to N_3_-kethoxal treatment to label ssDNA, after which the kethoxal is washed away. Next, cells are immediately processed through the standard ATAC-seq protocol, with Tn5 inserting into open chromatin regions. DNA is then isolated, and a click reaction is carried out, followed by streptavidin pull-down and library amplification. To control for sample loss and other potential biases associated with the pulldown procedure, we generated not just paired regular ATAC-seq libraries (from kethoxal-treated cells, but without a click reaction and biotin pulldown, in order to control for any effects of the kethoxal treatment), but also (for some key experiments) a “biotin-ATAC” control, in which no kethoxal labeling was carried out, but a transposase carrying biotinylated adapters was used, then these fragments were pulled down following the same procedure that was applied for KAS-ATAC (Figure 1b).

We carried out KAS-ATAC (together with parallel KAS-seq and ATAC-seq) in GM12878 lymphoblastoid cells and in HEK293 embryonic kidney cells as these have long been among the main cell lines of the ENCODE Project Consortium ^34,35^ and a wealth of other functional genomic data exists for them. Because we expected to recover less DNA than for regular ATAC libraries given that KAS-ATAC captures only a subset of the accessible genome and that the traditional ATAC-seq protocol only uses 50,000 cells as input, we pool at least two separate transposition reactions of 50,000 cells each for a given KAS-ATAC library.

As an initial assessment of the content of KAS-ATAC libraries, we examined KAS-ATAC, ATAC-seq and KAS-seq profiles over the mitochondrial genome (Figure 1c; Supplementary Figure 1); this was for two reasons: first, the wellknown high abundance of mitochondrial reads in ATAC libraries due to the absence of nucleosomes in the mitochondrial nucleoid, leaving it highly susceptible to Tn5 insertion ^12,36^, and second, that the mitochondrial genome contains the most abundant ssDNA structure in cells – the D-loop in its non-coding region ^37^. The D-loop is the major site of replication and transcription initiation in mitochondria ^38,39^ and exists as a triple-stranded structure, i.e. one of these three strands has numerous unpaired guanines and is accordingly highly enriched in KAS-seq libraries. We observe that while regular ATAC-seq libraries are slightly depleted over the D-loop, KAS-ATAC libraries are highly enriched over this region of chrM, with a broadly similar, but also distinct in some places profile compared to KAS-seq libraries. Thus, we conclude that KAS-ATAC indeed captures DNA fragments that are both accessible and single-stranded. Based on KAS-ATAC and ATAC-seq coverage over the D-loop, we estimate that we obtain at least 30*×* enrichment in KAS-ATAC over the transposition background. The overall fraction of reads originating from mitochondria was similar between all our matching KAS-ATAC and ATAC control libraries (Supplementary Figure 2).

Next, we examined the broad properties of KAS-ATAC datasets in the nuclear genome. Typical ATAC-seq libraries exhibit a characteristic periodic fragment length distribution ^12^, featuring a prominent subnucleosomal peak, corresponding to DNA fragments originating from within CREs, a second peak of size roughly that of the protection footprint of single nucleosomes in the immediate vicinity of CREs (as well as less frequent transposition events throughout the genome), a third dinucleosomal peak, and so on. We observe a similar picture in KAS-ATAC libraries (Figure 1d); however, while in ATAC-seq libraries the subnucleosomal peak is often (though not always) the strongest, all KAS-ATAC libraries we have examined feature relatively more prominent nucleosomal peaks (Supplementary Figure 3). This is to a certain extent counterintuitive and unexpected, given that the strongest KAS-seq peaks are those in the promoter regions of genes, corresponding to paused and initiating RNA polymerases; therefore, one may expect subnucleosomal fragments to dominate KAS-ATAC libraries. However, on the other hand much of ssDNA globally originates from transcribed intragenic regions, where nucleosomes are mostly intact. Indeed, we compared fragment length distributions across ENCODE chromatin states (annotated by the chromHMM algorithm ^40^), and we observed that the mononucleosomal peak is most prominent relative to the subnucleosomal one in the actively transcribed intragenic states (e.g. state 5 “Tx”; Supplementary Figures 5 and 4).

KAS-ATAC libraries also display a distinct signature around transcription start sites (TSSs), where a sharper peak is observed right at the TSS in metaprofile plots (Figure 1e; Supplementary Figure 6), likely corresponding to fragments associated with initiating/paused RNA polymerase.

### The single-stranded accessible chromatin landscape is distinct from total open chromatin genomic maps

We then surveyed local single-stranded accessible profiles around the genome. Figures 1f-g show representative KAS-seq, KAS-ATAC and ATAC-seq snapshots in GM12878 cells. Both ATAC-seq and KAS-ATAC exhibit enrichment over promoters and distal regulatory elements in the genome; however, in a number of cases KAS-ATAC peaks over promoter-distal CREs are considerably weaker than ATAC-seq signal over the same regions (e.g. highlighted areas in Figures 1f-g).

We quantified global differences between ATAC and KAS-ATAC by carrying out differential accessibility analysis using DESeq2 ^41^ on the set of common peaks called for both assays. In GM12878 cells, 1,948 and 4,641 regions were identified as preferentially enriched in ATAC-seq and KASATAC, respectively, and we identified 2,064 and 10,581 such peaks in HEK293 cells (Figure 2a-b). We compared these differential regions against chromHMM ^40^ chromatin state annotations for these two cell lines and found that they are not uniformly and randomly distributed across chromatin states (Figure 2c). KAS-ATAC libraries are relatively enriched over active TSSs (“TssA” state) and very strongly enriched over transcribed regions (“Tx” state), as well as over intragenic enhancer regions (“EnhG1/2”). They are depleted over active intergenic enhancers and over weak enhancers (“EnhG1” and “EnhWk”), and bivalent TSSs and enhancers. These observations generalized the anecdotal trend observed at individual loci – accessible intergenic enhancers appear to be less frequently engaged with polymerase bubbles than promoter regions. ATAC-seq and KAS-ATAC profiles around promoters and distal CREs also corroborate this conclusion (Figure 2d) – the two assays exhibit broadly similar profiles within promoters but in distal CREs it is often the case that KAS-ATAC signal is weaker.

**Figure 2:**
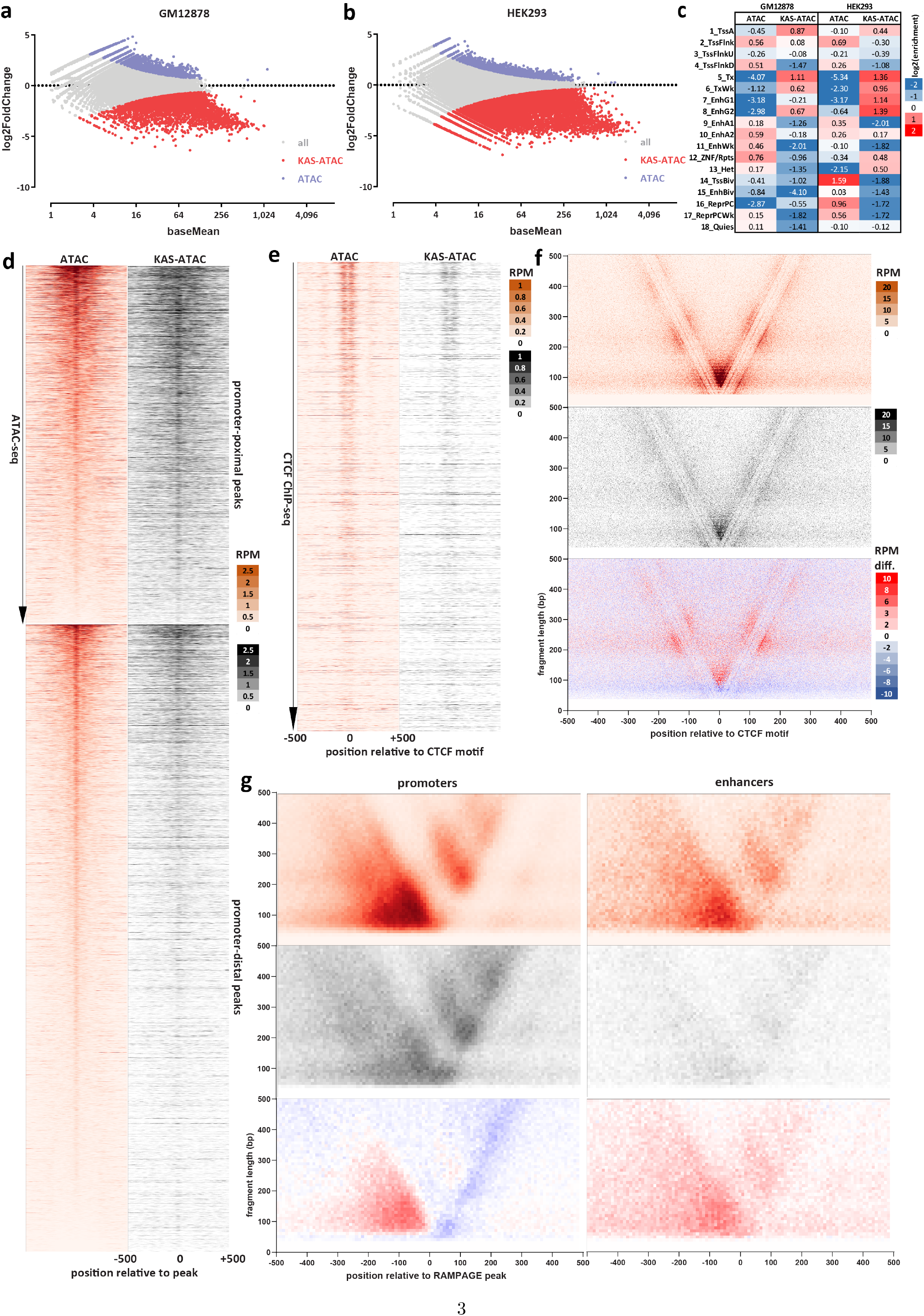
The single-stranded accessible chromatin landscape is distinct from total open chromatin genomic maps. (a) DESeq2 analysis of GM12878 ATAC and KAS-ATAC libraries. (b) DESeq2 analysis of HEK293 ATAC and KAS-ATAC libraries. (c) Enrichment of KAS-ATAC/ATAC differential peaks over chromHMM chromatin states. Enrichment is calculated as the *log*_2_ ratio of the observed versus expected fractions of peaks overlapping each state (based on the overlap of the total set of peaks with chromHMM states). (d) ATAC-seq and KAS-ATAC profiles around promoter-proximal and distal peaks (GM12878 cells). (e) ATAC-seq and KAS-ATAC profiles around occupied CTCF sites in GM12878 cells (excluding promoter-proximal CTCF peaks; CTCF ChIP-seq peak calls were obtained from ENCODE accession ENCFF827JRI). (f) ATAC-seq, KAS-ATAC, and KAS-ATAC/ATAC differential V-plot ^25^ around occupied CTCF sites. (g) ATAC-seq, KAS-ATAC, and KAS-ATAC/ATAC differential V-plot ^25^ around RAMPAGE ^26^ peaks in promoter and in distal CRE regions (RAMPAGE peaks were obtained from ENCODE accession ID ENCSR000AEI).

Next, we turned our attention to the properties of sites occupied by the CTCF protein, which is the main insulator factor in mammalian cells ^42^. We excluded promoter-proximal CTCF sites from our analysis and focused on distal CTCF peaks only. KAS-ATAC signal is notably weaker at CTCF sites than ATAC-seq (Figure 2e). CTCF also has a very strong effect on nucleosome positioning through its very robust and persistent occupancy of its own binding sites, often fixing the positions of more than a dozen nucleosomes around it ^43^; CTCF sites thus display a characteristic V-plot ^25^ signature in ATAC-seq datasets, with a strong subnucleosomal footprint corresponding to the binding of CTCF itself and larger fragments including neighboring nucleosomes surrounding it (Figure 2f). This signature is also observed in KAS-ATAC data, although with a reduced magnitude. However, it has been previously reported that G-quadruplexes promote CTCF binding in mammalian cells ^44^, and this could account for our observations of relatively weak KAS-ATAC signal in these datasets at CTCF sites. Alternatively, this signal might be the product of background DNA “breathing” or DNA replication intermediates that CTCF associates with, as there is only modest enrichment of CTCF sites in regular KAS-seq libraries (Supplementary Figure 7).

We also analyzed fragment structure around transcription initiation sites. To this end, we took advantage of existing ENCODE TSS annotations for total RNA obtained from RAMPAGE ^26^ datasets, which we divided into promoter-proximal and promoter-distal groups, and then analyzed ATAC and KAS-ATAC fragment distributions (in a transcription directionality-aware manner; Figure 2g). ATAC-seq V-plots around promoters feature a prominent mass of subnucleosomal and some mononucleosomal fragments immediately upstream of the TSS (corresponding to occupancy by regulatory factors, and general transcription factors), another group of subnucleosomal fragments immediately around the TSS (likely corresponding to RNA polymerase initiation complexes), as well as +1 nucleosome and dinucleosomal-fragment masses downstream of the TSS. This signature is preserved in KAS-ATAC, but with important differences – the upstream subnucleosomal fragments are depleted in KAS-ATAC, while the initiation complex ones are enriched, as is the +1 nucleosome and the downstream dinucleosome. While transcription initiation in mammals has long been demonstrated to be bidirectional ^28,45^, these results suggest preferential absolute abundance of polymerase bubbles in the primary direction of transcription. In addition, engaged polymerase at the promoter and immediately downstream of it is associated with intact nucleosomes.

In contrast to promoters, transcribed distal CREs exhibit a similar picture in its broad structure in ATAC-seq data, but this signature is very weak in KAS-ATAC, and there is no preferential enrichment of KAS-ATAC fragments downstream of the transcription initiation site.

### Estimating the absolute abundance of accessible ssDNA

KAS-ATAC also offers an opportunity to address the question of how often accessible DNA fragments contain ssDNA inside them in absolute terms. To answer this question, we used a paired biotin-ATAC dataset so that we could control as closely as possible for the inevitable loss of material during sample handling and to obtain comparable in their absolute molecular content ATAC and KAS-ATAC libraries. To maximize sample recovery, we also combined 12 normal ATAC samples (of 50,000 cells each, for a total of 600,000 cells input) into each of these libraries, which were generated in HEK293 cells. We sequenced them to saturation, and estimated the total library complexity using the `preseq` algorithm ^46^ (Figure 3a). As expected, the total molecular complexity of KAS-ATAC libraries is lower than that of ATAC-seq ones, by *∼*50%.

**Figure 3:**
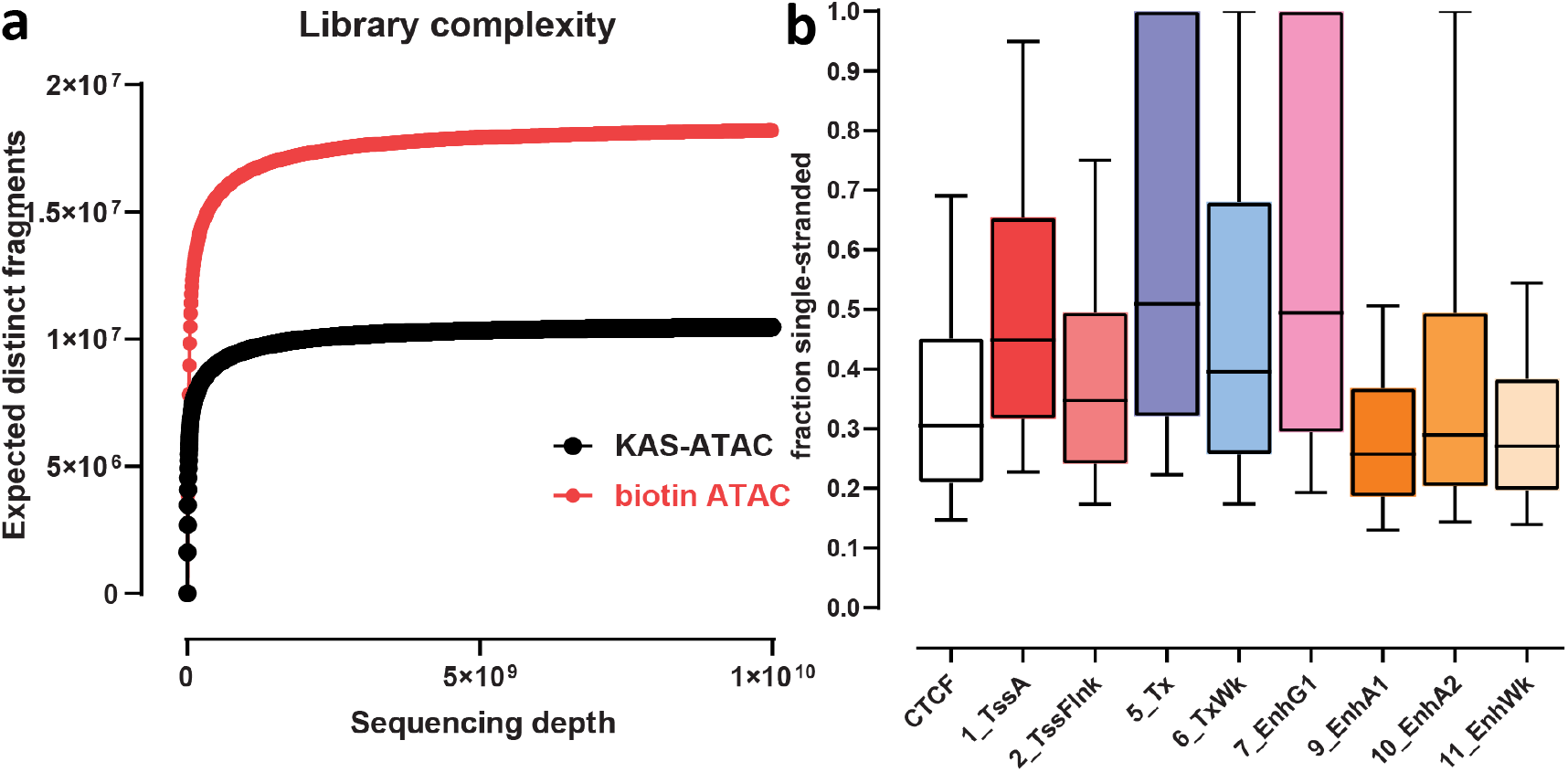
Estimating absolute levels of single-stranded accessible DNA. (a) Estimates for total library complexity in KAS-ATAC and biotin-ATAC HEK293 samples. (b) Distribution of the ratios between the number of KAS-ATAC molecules and biotin-ATAC molecules for ATAC-seq peaks found in different chromatin states and for CTCF occupancy sites.

We used the total molecular complexity of the KASATAC and biotin-ATAC samples to calculate the total number of fragments (that was derived from *∼*600,000 cells) for each ATAC-seq peak found in a given chromHMMM state as well as for CTCF peaks, and then estimated the ratio between the two to evaluate how often accessible chromatin regions are also associated with ssDNA (Figure 3b; Supplementary Figure 8). The median estimate we obtain for active TSSs is that *∼*45% of ATAC fragments contain ssDNA. The highest such estimates are for peaks in the actively transcribed (“Tx”) and intragenic enhancer (“EnhG1”) states, at *∼*50%, but also often approaching 100%. In contrast, for active intergenic enhancers the median fraction of ssDNA fragments is in the 20-25% range (“EnhA1/A2”). For CTCF sites the median estimate is *∼*30%. These results corroborate the observation of lower levels of transcription at intergenic enhancers, and of the presence of ssDNA structures at CTCF sites, while also quantifying their abundance in cells.

### Transcription factor footprints in KAS-ATAC

Finally, we asked whether transcription factors are associated with DNA in a similar way regardless of whether transcription (or other processes generating ssDNA) are occurring in the immediate vicinity. To this end, we calculated aggregate transcription factor footprints over TF motifs within ChIP-seq peaks for each TF in GM12878 and HEK293 cells, including all such TFs for which ChIPseq data is available from the ENCODE Project Consortium ^34^ and for which motifs are annotated in the CISBP database ^47^. We applied bias-correction utilizing bias models trained using the ChromBPNet framework (see the Methods section for detail), as Tn5 insertion bias is a well-documented confounding issue in ATAC-seq datasets ^48–52^.

For most TFs we find similar depth of footprints in both ATAC and KAS-ATAC datasets (Figure 4a-d; Supplementary Figures 9–12 For example, CTCF footprints in HEK293 cells are largely the same in both the total ATAC library and within ssDNA-containing fragments (Figure 4a), with concordant footprint depths and fine-scale features. Analogous conclusions apply to factors such as the bZIP family factor ATF3 (Figure 4b), the nuclear receptor NR2F1 (Figure 4c), the YY1 zing finger TF (Figure 4d), and most others (Supplementary Figures 9–12). Thus, most transcription factors appear to physically associate with DNA fragments where active transcription is ongoing in the immediate vicinity.

**Figure 4:**
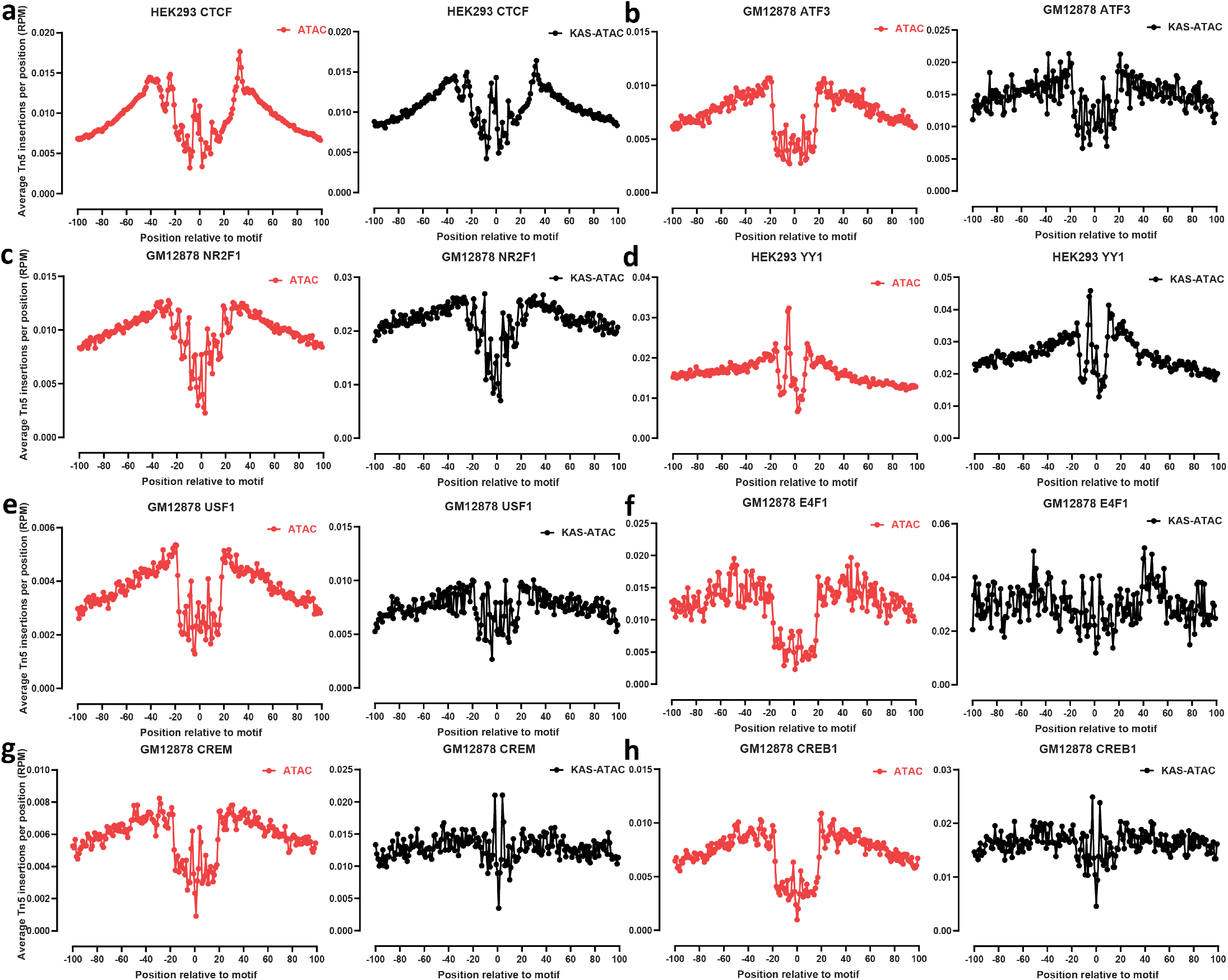
Transcription factor footprints in KAS-ATAC and ATAC datasets. Shown are the aggregate average Tn5 insertion profiles (see Methods for details) around the motifs for the indicated factors that are located within ENCODE ChIP-seq peaks for the TF in the given cell line. (a) HEK293 CTCF; (b) GM12878 ATF3; (c) GM12878 NR2F1; (d) HEK293 YY1; (e) GM12878 USF1; (f) GM12878 E4F1; (g) GM12878 CREM; (h) GM12878 CREB1.

However, we also find several TFs for which strong footprints are only observed in ATAC-seq libraries and not in KAS-ATAC. Examples include the bHLH TF USF1 (Figure 4e), the zinc finger E4F1 factor (Figure 4f), and the bZIP family TFs CREM and CREB1 (Figure 4g-h). These are factors which appear to preferentially associate with DNA fragments that are not actively undergoing transcription, although the precise mechanisms underlying this phenomenon are still unclear for each individual case.

## Summary

In this work, we describe the KAS-ATAC assay, which we developed to identify the subset of physically accessible DNA fragments in the genome that also contain embedded ssDNA structures, the bulk of which is the result of the opening of the DNA double strand associated with engaged RNA polymerase molecules. We use KAS-ATAC to answer several questions regarding the relationship between active transcription/ssDNA formation and fine-scale chromatin architecture.

We show that the genomic regions that are jointly accessible and containing ssDNA are preferentially located in promoters and inside gene bodies. In comparison, distal enhancers, although often comparably accessible to promoters, are considerably less frequently transcribed. These two classes of elements are also characterized by subtle differences in their subnucleosomal and nucleosomal organization as it relates to the act of transcription.

We evaluate the absolute fraction of accessible DNA that is engaged in transcription for different classes of regulatory elements, estimating that up to half of fragments within promoters contain ssDNA, while in distal enhancers this fraction is less than a quarter of fragments.

Finally, we demonstrate that transcription factors physically associate with actively transcribed DNA, and most of them do so with comparable frequency to non-ssDNA-containing DNA. However, a number of TFs are in fact depleted in KAS-ATAC libraries and are preferentially associated with non-ssDNA DNA fragments.

We expect KAS-ATAC to be a useful tool for further future exploration of the detailed relationship between transcription/ssDNA structures and the physical organization of chromatin in many other context, as will the combination of kethoxal labeling and other molecular readouts of the chromatin structure.

## Methods

### Cell culture

GM12878 and HEK293T cells were grown using the standard protocols for each cell lines – RPMI 1640, 2 mM L-glutamine, 15% fetal bovine serum and 1% penicillinstreptomycin for GM12878 cells, and DMEM + sodium pyruvate, 6 mM L-glutamine, 10% fetal bovine serum and 1% penicillin-streptomycin for HEK293T cells.

### KAS-seq

KAS-seq experiments were executed as previously described ^53^.

For HEK293 cells, media was removed and roomtemperature 1*×* PBS was used to wash them, then they were dissociated with trypsin, which was quenched with media. Cells were pelleted at room temperature, and then resuspended in 500 *µ*L of media supplemented with 5 mM N_3_-kethoxal (final concentration). GM12878 cells were pelleted down by centrifugation at room temperature and also resuspended in 500 *µ*L of media supplemented with 5 mM N_3_-kethoxal (final concentration). Cells were incubated for 10 minutes at 37 °C with shaking at 500 rpm in a Thermomixer, then pelleted by centrifuging at 500 *g* for 5 minutes at 4 °C. Genomic DNA was extracted using the Monarch gDNA Purification Kit (NEB T3010S) following the standard protocol but with elution using 175 *µ*L 25 mM K_3_BO_3_ at pH 7.0.

The click reaction was carried out by combining transposed DNA, 5 *µ*L 20 mM DBCO-PEG4-biotin (DMSO solution, Sigma 760749), 20 mM K_3_BO_3_ (pH 7.0) and 20 *µ*L 10*×* PBS in a final volume of 200 *µ*L.

Biotinylated DNA was purified using AMPure XP beads and sheared on a Covaris E220 instrument down to *∼*150-200 bp size.

For streptavidin pulldown of biotin-labeled DNA, 10 *µ*L of 10 mg/mL Dynabeads MyOne Streptavidin T1 beads (Life Technologies, 65602) were separated on a magnetic stand, then washed with 300 *µ*L of 1*×* TWB (Tween Washing Buffer; 5 mM Tris-HCl pH 7.5; 0.5 mM EDTA; 1 M NaCl; 0.05% Tween 20). The beads were resuspended in 300 *µ*L of 2*×* Binding Buffer (10 mM Tris-HCl pH 7.5, 1 mM EDTA; 2 M NaCl), the sonicated DNA was added (diluted to a final volume of 300 *µ*L if necessary), and the beads were incubated for *≥*15 minutes at room temperature on a rotator. After separation on a magnetic stand, the beads were washed with 300 *µ*L of 1*×* TWB, and heated at 55 °C in a Thermomixer with shaking for 2 minutes. After removal of the supernatant on a magnetic stand, the TWB wash and 55 °C incubation were repeated.

Final libraries were prepared on beads using the NEBNext Ultra II DNA Library Prep Kit (NEB, #E7645) as follows. End repair was carried out by resuspending beads in 50 *µ*L 1*×* EB buffer, and adding 3 *µ*L NEB Ultra End Repair Enzyme and 7 *µ*L NEB Ultra End Repair Enzyme, followed by incubation at 20 °C for 30 minutes (in a Thermomixer, with shaking at 1,000 rpm) and then at 65 °C for 30 minutes.

Adapters were ligated to DNA fragments by adding 30 *µ*L Blunt Ligation mix, 1 *µ*L Ligation Enhancer and 2.5 *µ*L NEB Adapter, incubating at 20 °C for 20 minutes, adding 3 *µ*L USER enzyme, and incubating at 37 °C for 15 minutes (in a Thermomixer, with shaking at 1,000 rpm) .

Beads were then separated on a magnetic stand, and washed with 300 *µ*L TWB for 2 minutes at 55 °C, 1000 rpm in a Thermomixer. After separation on a magnetic stand, beads were washed in 100 *µ*L 0.1 *×* TE buffer, then resuspended in 15 *µ*L 0.1 *×* TE buffer, and heated at 98 °C for 10 minutes.

For PCR, 5 *µ*L of each of the i5 and i7 NEB Next sequencing adapters were added together with 25 *µ*L 2*×* NEB Ultra PCR Mater Mix. PCR was carried out with a 98 °C incubation for 30 seconds and 12 cycles of 98 °C for 10 seconds, 65 °C for 30 seconds, and 72 °C for 1 minute, followed by incubation at 72 °C for 5 minutes.

Beads were separated on a magnetic stand, and the supernatant was cleaned up using 1.8*×* AMPure XP beads.

Libraries were sequenced in a paired-end format on an Illumina NextSeq instrument using NextSeq 500/550 high output kits (2*×*38 cycles).

### Tn5 production

Tn5 transposase was produced as previously described ^54^, with the following modifications of the adapter oligos for biotin-ATAC experiments:

Tn5MErev:

`/5Phos/CTGTCTCTTATACACATCT`

Tn5ME-A:

`/5Biotin/TCGTCGGCAGCGTCAGATGTGTATAAGAGACAG`

Tn5ME-B:

`/5Biotin/GTCTCGTGGGCTCGGAGATGTGTATAAGAGACAG`

### ATAC-seq experiments

ATAC-seq experiments were carried out following the Omni-ATAC protocol as previously described ^55,56^. Briefly, *∼*50,000 cells (with or without prior kethoxal treatment) were centrifuged at 500 *g*, then resuspended in 500 *µ*L 1*×* PBS and centrifuged again. Cells were then resuspended in 50 *µ*L ATAC-RSB-Lysis buffer (10 mM Tris-HCl pH 7.4, 10 mM NaCl, 3 mM MgCl_2_, 0.1% IGEPAL CA-630, 0.1% Tween-20, 0.01% Digitonin) and incubated on ice for 3 minutes. Subsequently 1 mL ATAC-RSB-Wash buffer (10 mM Tris-HCl pH 7.4, 10 mM NaCl, 3 mM MgCl_2_, 0.1% Tween20, 0.01% Digitonin) were added, the tubes were inverted several times, and nuclei were centrifuged at 500 *g* for 5 min at 4 °C.

Transposition was carried out by resuspending nuclei in a mix of 25 *µ*L 2*×* TD buffer (20 mM Tris-HCl pH 7.6, 10 mM MgCl_2_, 20% Dimethyl Formamide), 2.5 *µ*L Tn5 transposase (custom produced) and 22.5 *µ*L nuclease-free H_2_O, and incubating at 37 °C for 30 min in a Thermomixer at 1000 RPM.

Transposed DNA was isolated using the MinElute PCR Purification Kit (Qiagen Cat# 28004/28006), and PCR amplified as previously described ^55,56^ (72 °C for 3 minutes, 98 °C for 30 seconds, 10 cycles of 98 °C for 10 seconds, 63 °C for 30 seconds, 72 °C for 30 seconds). Libraries were purified using the MinElute kit, then sequenced on an Illumina NextSeq 550 as 2*×*38mers.

### KAS-ATAC experiments

For KAS-ATAC experiments, cells were processed and subjected to N_3_-kethoxal treatment as described above for KAS-seq.

Subsequently they were washed by centrifugation and removal of media, resuspension in 500 *µ*L 1*×* PBS, and centrifugation. The ATAC-seq step was then carried out, by resuspending cells in 50 *µ*L ATAC-RSB-Lysis buffer, incubation on ice for 3 minutes, addition of 1 mL ATAC-RSB-Wash buffer, centrifugation at 500 *g* for 5 min at 4 °C, and transposition in a total volume of 50 *µ*L. DNA purification was carried out using the MinElute PCR Purification Kit. Multiple parallel ATAC reactions were carried out, including *∼*50,000 cells each, then were pooled together for the click reaction and biotin pull down.

Click reaction was carried out as described above, by combining 175 *µ*L transposed DNA, 5 *µ*L 20 mM DBCOPEG4-biotin (DMSO solution, Sigma 760749), 20 mM K_3_BO_3_ (pH 7.0) and 20 *µ*L 10*×* PBS in a final volume of 200 *µ*L. Biotinylated DNA was purified using the MinElute PCR Purification Kit.

For streptavidin pulldown of biotin-labeled DNA, 10 *µ*L of 10 mg/mL Dynabeads MyOne Streptavidin T1 beads (Life Technologies, 65602) were separated on a magnetic stand, then washed with 300 *µ*L of 1*×* TWB (Tween Washing Buffer; 5 mM Tris-HCl pH 7.5; 0.5 mM EDTA; 1 M NaCl; 0.05% Tween 20). The beads were resuspended in 300 *µ*L of 2*×* Binding Buffer (10 mM Tris-HCl pH 7.5, 1 mM EDTA; 2 M NaCl), the biotinylated DNA was added (diluted to a final volume of 300 *µ*L if necessary), and the beads were incubated for *≥*15 minutes at room temperature on a rotator. After separation on a magnetic stand, the beads were washed with 300 *µ*L of 1*×* TWB, and heated at 55 °C in a Thermomixer with shaking for 2 minutes. After removal of the supernatant on a magnetic stand, the TWB wash and 55 °C incubation were repeated.

Libraries were prepared on beads using the same settings as described above (72 °C for 3 minutes, 98 °C for 30 seconds, 10 cycles of 98 °C for 10 seconds, 63 °C for 30 seconds, 72 °C for 30 seconds). Beads were separated on a magnetic stand, and the supernatant was cleaned up using the MinElute PCR Purification Kit.

### Biotin-ATAC experiments

ATAC-seq experiments were carried out as described above but using a custom-produced biotinylated Tn5 transposase to tagment chromatin.

Purified transposed DNA was then subjected to streptavidin pulldown and PCR amplification on beads as described above.

### ATAC-seq, KAS-seq, and KAS-ATAC data processing

Computational processing was carried out as previously described ^53,57^

Demultipexed FASTQ files were mapped to the `hg38` assembly of the human genome as 2*×*36mers using Bowtie ^58^ with the following settings: `-v 2 -k 2 -m 1 --best`

`--strata`. Mitochondrial mapping reads were filtered out and duplicate reads were removed using `picard-tools` (version 1.99).

Reads were also mapped separately to the mitochondrial genome using the following settings: `-v 2 -a --best--strata`, both in order to estimate the extent of mitochondrial contamination and to generate mitochondrial genome coverage tracks.

Browser tracks generation, fragment length estimation, TSS enrichment calculations, and other analyses were carried out using custom-written Python scripts https://github.com/georgimarinov/GeorgiScripts. The refSeq annotation was used for evaluation of enrichment around TSSs.

Peak calling was carried out using MACS2 ^59^, with the following settings: `-g hs -f BAMPE --to-large --keep-dup all --nomodel`.

Absolute library complexity was estimated using the `lc_extrap` function in the `preseq` package ^46^.

For differential accessibility analysis, a set of unified peaks was compiled across all libraries and conditions being compared, and read counts were calculated for each peak. Differential analysis was the carried out using DESeq2 ^41^.

### Transcription factor motif analyses

As a first step, motif locations for all human TF motifs in the CIS-BP database ^47^ were identified genome-wide using FIMO ^60^. These maps were then intersected with ChIP-seq peaks from the ENCODE Project Consortium (ENCODE IDs: GM12878 ARID3A: ENCFF531CLQ; GM12878 ARNT: ENCFF596RIT; GM12878 ATF3: ENCFF882AEU; GM12878 ATF7: ENCFF277FJJ; GM12878 BACH1: ENCFF631RZH; GM12878 BATF: ENCFF728KFD; GM12878 BCL11A: ENCFF824QXX; GM12878 BHLHE40: ENCFF445XYV; GM12878 BRCA1: ENCFF303GOQ; GM12878 CBFB: ENCFF876CUT; GM12878 CEBPB: ENCFF864EON; GM12878 CEBPZ: ENCFF639KLW; GM12878 CREB1: ENCFF053UZX; GM12878 CREM: ENCFF294IGP; GM12878 CTCF: ENCFF796WRU; GM12878 CUX1: ENCFF451AII; GM12878 E2F4: ENCFF744QAC; GM12878 E2F8: ENCFF961ZFW; GM12878 E4F1: ENCFF855PXT; GM12878 EBF1: ENCFF895MHN; GM12878 EGR1: ENCFF612EIU; GM12878 ELF1: ENCFF146SYU; GM12878 ELK1: ENCFF164MPE; GM12878 ESRRA: ENCFF660PIB; GM12878 ETS1: ENCFF568AZT; GM12878 ETV6: ENCFF151UJT; GM12878 FOS: ENCFF571DGT; GM12878 FOXM1: ENCFF549GKZ; GM12878 GABPA: ENCFF093KLR; GM12878 HSF1: ENCFF663ODL; GM12878 IRF3: ENCFF785MSW; GM12878 IRF4: ENCFF113VGD; GM12878 IRF5: ENCFF478SRO; GM12878 JUNB: ENCFF912OPT; GM12878 JUND: ENCFF023QWA; GM12878 KLF5: ENCFF992IBT; GM12878 MAFK: ENCFF436SJS; GM12878 MAX: ENCFF361EVH; GM12878 MAZ: ENCFF250FJC; GM12878 MEF2A: ENCFF826GQU; GM12878 MEF2B: ENCFF884QQW; GM12878 MEF2C: ENCFF238UKB; GM12878 MXI1: ENCFF376AEL; GM12878 MYB: ENCFF705CGM; GM12878 MYC: ENCFF214XPD; GM12878 NFATC1: ENCFF172KBM; GM12878 NFATC3: ENCFF002XEC; GM12878 NFYA: ENCFF937QTP; GM12878 NFYB: ENCFF156MUM; GM12878 NR2C1: ENCFF626EEU; GM12878 NR2C2: ENCFF782KRV; GM12878 NR2F1: ENCFF118UKC; GM12878 NRF1: ENCFF864EBU; GM12878 PAX5: ENCFF192WNJ; GM12878 PAX8: ENCFF934NVD; GM12878 PBX3: ENCFF402DQD; GM12878 PKNOX1: ENCFF946JFA; GM12878 POU2F2: ENCFF934JFA; GM12878 RELA: ENCFF141SMI; GM12878 RELB: ENCFF968MBN; GM12878 REST: ENCFF262MRD; GM12878 RFX5: ENCFF601WHE; GM12878 RUNX3: ENCFF346JDW; GM12878 RXRA: ENCFF063KLF; GM12878 SIX5: ENCFF878FBM; GM12878 SMAD1: ENCFF033MGN; GM12878 SP1: ENCFF038AVV; GM12878 SPI1: ENCFF492ZRZ; GM12878 SREBF1: ENCFF397YRO; GM12878 SREBF2: ENCFF896MPE; GM12878 SRF: ENCFF089YRX; GM12878 STAT1: ENCFF896SEI; GM12878 STAT3: ENCFF029PAA; GM12878 STAT5A: ENCFF006GSY; GM12878 TBX21: ENCFF170IZO; GM12878 TCF12: ENCFF068YYR; GM12878 TCF3: ENCFF700TAS; GM12878 TCF7: ENCFF900GTF; GM12878 USF1: ENCFF879TPT; GM12878 USF2: ENCFF196JQW; GM12878 YBX1: ENCFF488IGF; GM12878 YY1: ENCFF258UEW; GM12878 ZBED1: ENCFF753UAX; GM12878 ZBTB33: ENCFF845VSZ; GM12878 ZBTB4: ENCFF880BEK; GM12878 ZEB1: ENCFF703XCL; GM12878 ZNF274: ENCFF034UEJ; GM12878 ZNF384: ENCFF955FRU; HEK293 BCL11A: ENCFF972WFM; HEK293 BCL6B: ENCFF884UOX; HEK293 CTCF: ENCFF071FSR; HEK293 EGR2: ENCFF501WRO; HEK293 ELK4: ENCFF226QHL; HEK293 GFI1B: ENCFF199YAZ; HEK293 GLI2: ENCFF576SDC; HEK293 GLIS1: ENCFF628APG; HEK293 GLIS2: ENCFF311QUH; HEK293 HIC1: ENCFF979QGX; HEK293 HOXA7: ENCFF521UKW; HEK293 HOXB7: ENCFF610SSK; HEK293 HOXC10: ENCFF722GPK; HEK293 HOXD13: ENCFF223BOC; HEK293 KLF16: ENCFF585OJO; HEK293 KLF1: ENCFF103JPK; HEK293 KLF8: ENCFF790TTV; HEK293 MAZ: ENCFF529XTF; HEK293 MEIS1: ENCFF628KKY; HEK293 MZF1: ENCFF669FUB; HEK293 PBX3: ENCFF851ZEB; HEK293 PRDM1: ENCFF753PSU; HEK293 PRDM4: ENCFF351OYP; HEK293 REST: ENCFF437IAR; HEK293 SCRT1: ENCFF424QVD; HEK293 SCRT2: ENCFF916ITG; HEK293 SP2: ENCFF444VWF; HEK293 SP3: ENCFF663PCO; HEK293 TCF7L2: ENCFF136SOA; HEK293 WT1: ENCFF006ZKN; HEK293 YY1: ENCFF430PDB; HEK293 YY2: ENCFF755NLD; HEK293 ZBTB49: ENCFF066ZQN; HEK293 ZBTB6: ENCFF139IGE; HEK293 ZBTB7A: ENCFF114DGV; HEK293 ZEB1: ENCFF079WZH; HEK293 ZFHX2: ENCFF153XOB; HEK293 ZIC2: ENCFF359GIY; HEK293 ZNF148: ENCFF410SFG; HEK293 ZNF263: ENCFF108NXV; HEK293 ZNF274: ENCFF315BAF; HEK293 ZNF350: ENCFF064ZJO; HEK293 ZNF423: ENCFF222HPT; HEK293 ZSCAN16: ENCFF180POM; HEK293 ZSCAN4: ENCFF952OOU).

Aggregate footprints were generated for each TF by calculating the average number of Tn5 insertions (shifted by +4/-4 bp) normalized to RPM (Read Per Million). Bias correction was carried out using a genome-wide bias prediction track for a ChromBPNet model (https://github.com/kundajelab/chrombpnet) trained on the very deeply sequenced main ENCODE ATAC-seq dataset for GM12878 cells by subtracting the rescaled predicted bias profile from the observed ATAC/KAS-ATAC insertion profile.

## Data availability

Sequencing reads for libraries included in this manuscript have been submitted to the Gene Expression Omnibus and are available under GEO accession GSE264534.

## Acknowledgments

This work was supported by NIH grants (P50HG007735, RO1 HG008140, U19AI057266, UM1HG009442 and 1UM1HG009436 to W.J.G., the Rita Allen Foundation (to W.J.G.), the Baxter Foundation Faculty Scholar Grant, and the Human Frontiers Science Program grant RGY006S (to W.J.G). W.J.G. also acknowledges support by grants 2017-174468 and 2018-182817 from the Chan Zuckerberg Initiative. S.H.K. was supported by MSTP training grant T32GM007365 and the Paul and Daisy Soros Fellowship. Fellowship support was also provided by the Stanford School of Medicine Dean’s Fellowship (G.K.M.).

The authors would like to thank Anusri Pampari, Zohar Shipony, and Anna Shcherbina for technical assistance, and members of the Greenleaf, and Kundaje labs for helpful discussions and suggestions.

## Author contributions

G.K.M. and S.H.K. conceived the project. S.H.K. and G.K.M. carried out experiments. G.K.M. analyzed data. G.K.M. and S.H.K. wrote the manuscript with input from all authors.

## Competing interests

The authors declare no competing interests.

## Supplementary Materials

### Supplementary Figures

**Supplementary Figure 1:**
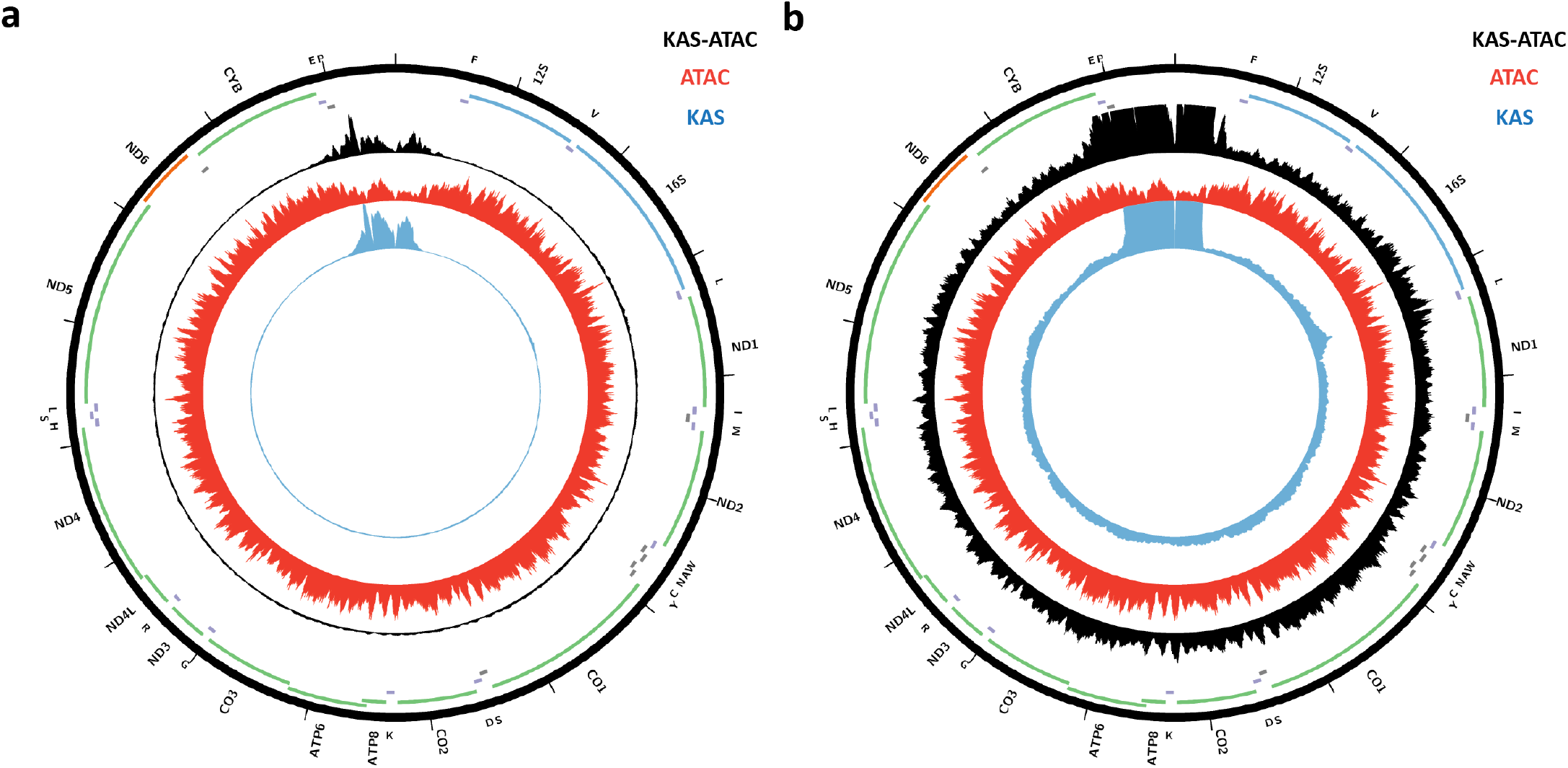
KAS-seq, ATAC-seq, and KAS-ATAC mitochondrial genome profiles in human GM12878 cells. (a) Shown are KAS-seq, ATAC-seq, and KAS-ATAC mitochondrial genome profiles (b) Shown are KAS-seq, ATAC-seq, and KAS-ATAC mitochondrial genome profiles with signal capped so that differences outside the D-loop can be visualized.

**Supplementary Figure 2:**
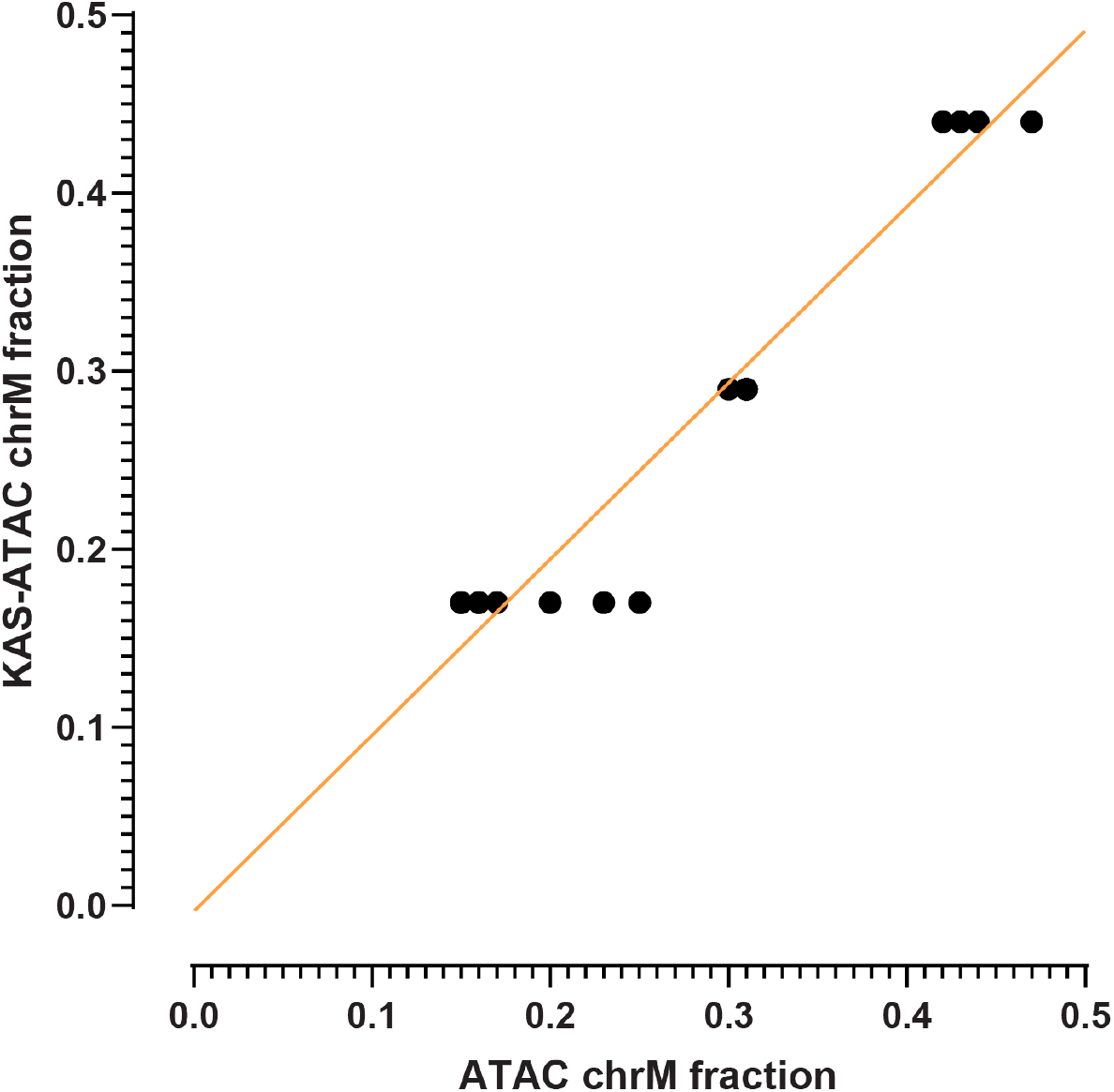
Fraction of mitochondrial reads in ATAC-seq and KAS-ATAC libraries.

**Supplementary Figure 3:**
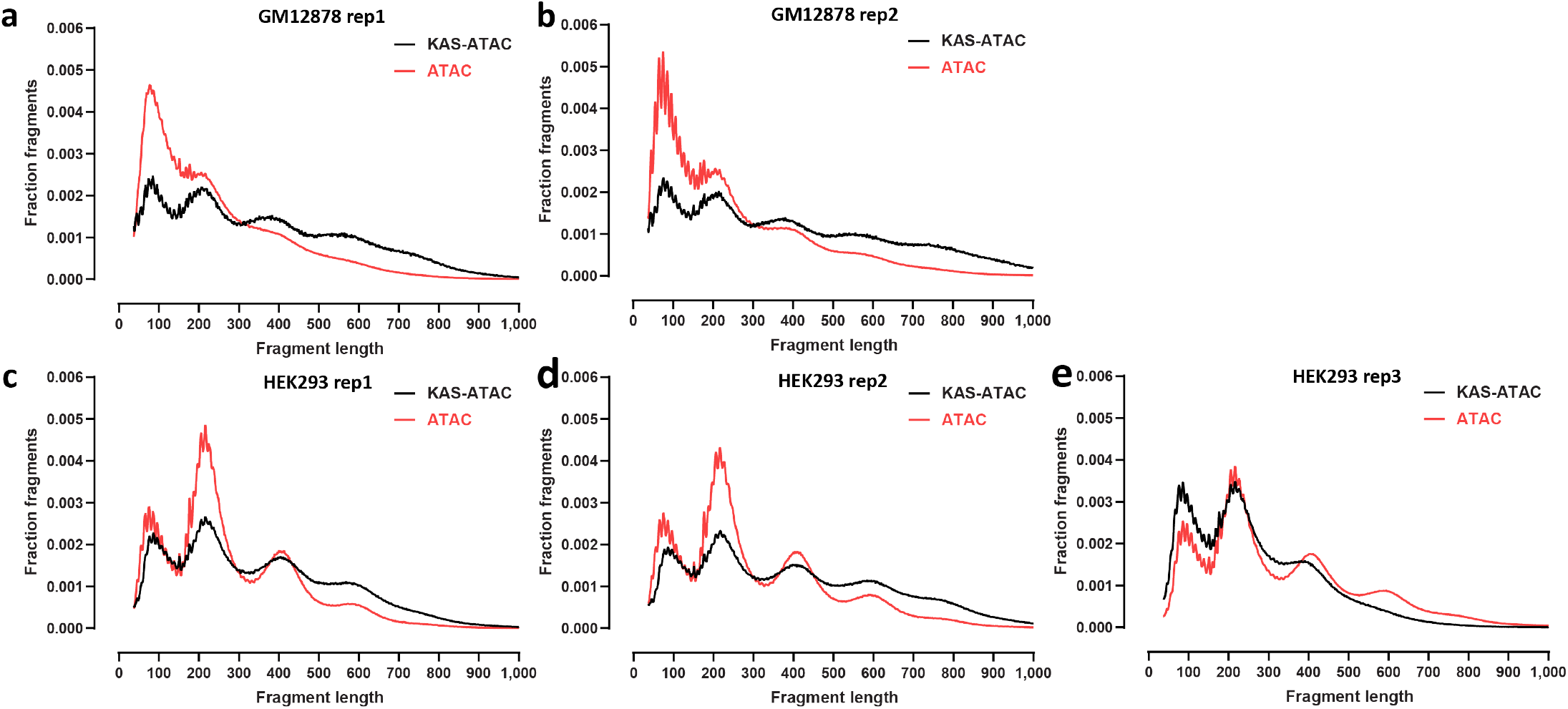
Fragment length distribution in ATAC-seq and KAS-ATAC. (a) GM12878 cells, replicate 1; (b) GM12878 cells, replicate 2; (c) HEK293 cells, replicate 1; (d) HEK293 cells, replicate 2; (e) HEK293 cells, replicate 3.

**Supplementary Figure 4:**
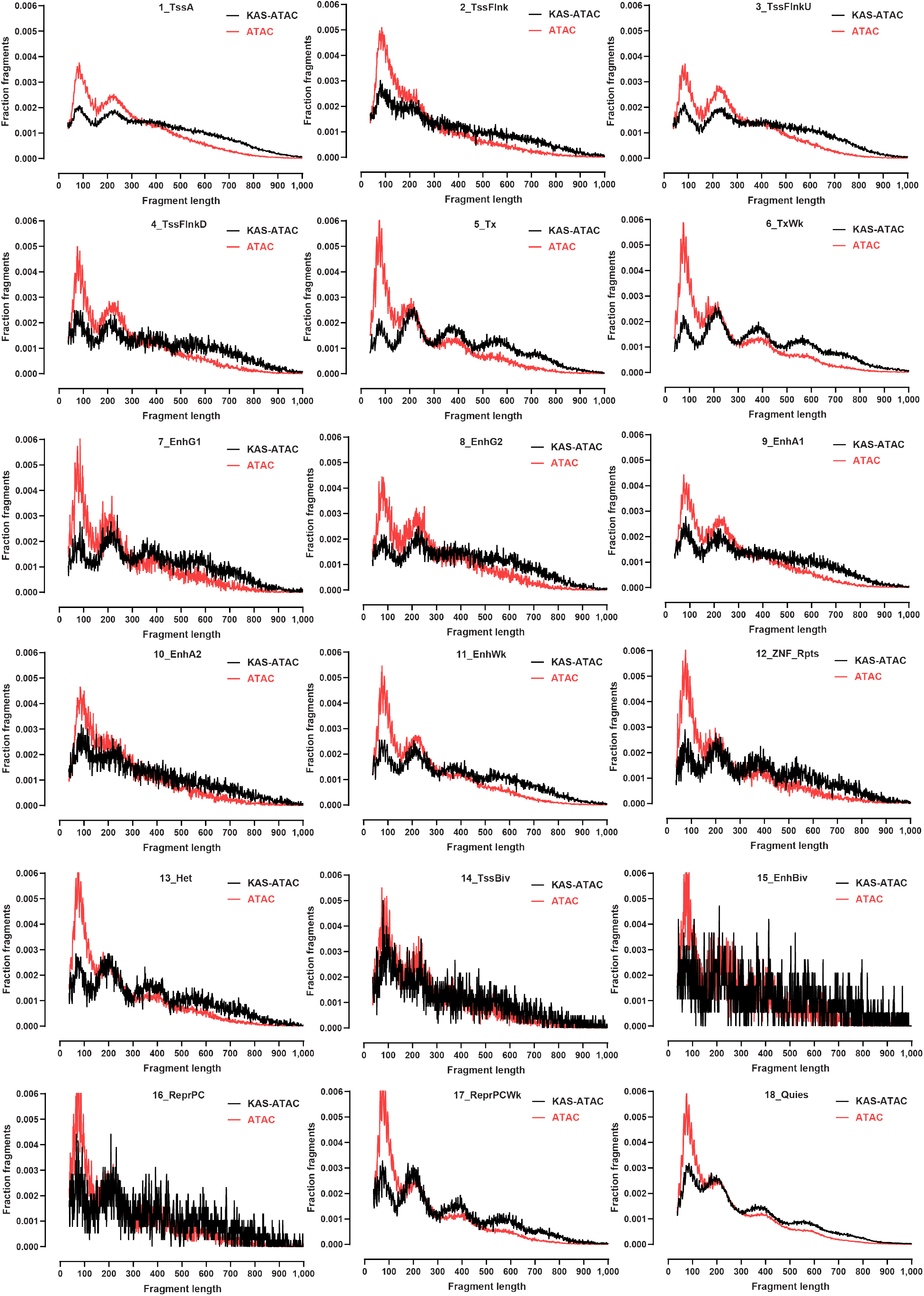
Distribution of KAS-ATAC and ATAC-seq fragment lengths in different chromatin states in GM12878 (replicate 1) cells.

**Supplementary Figure 5:**
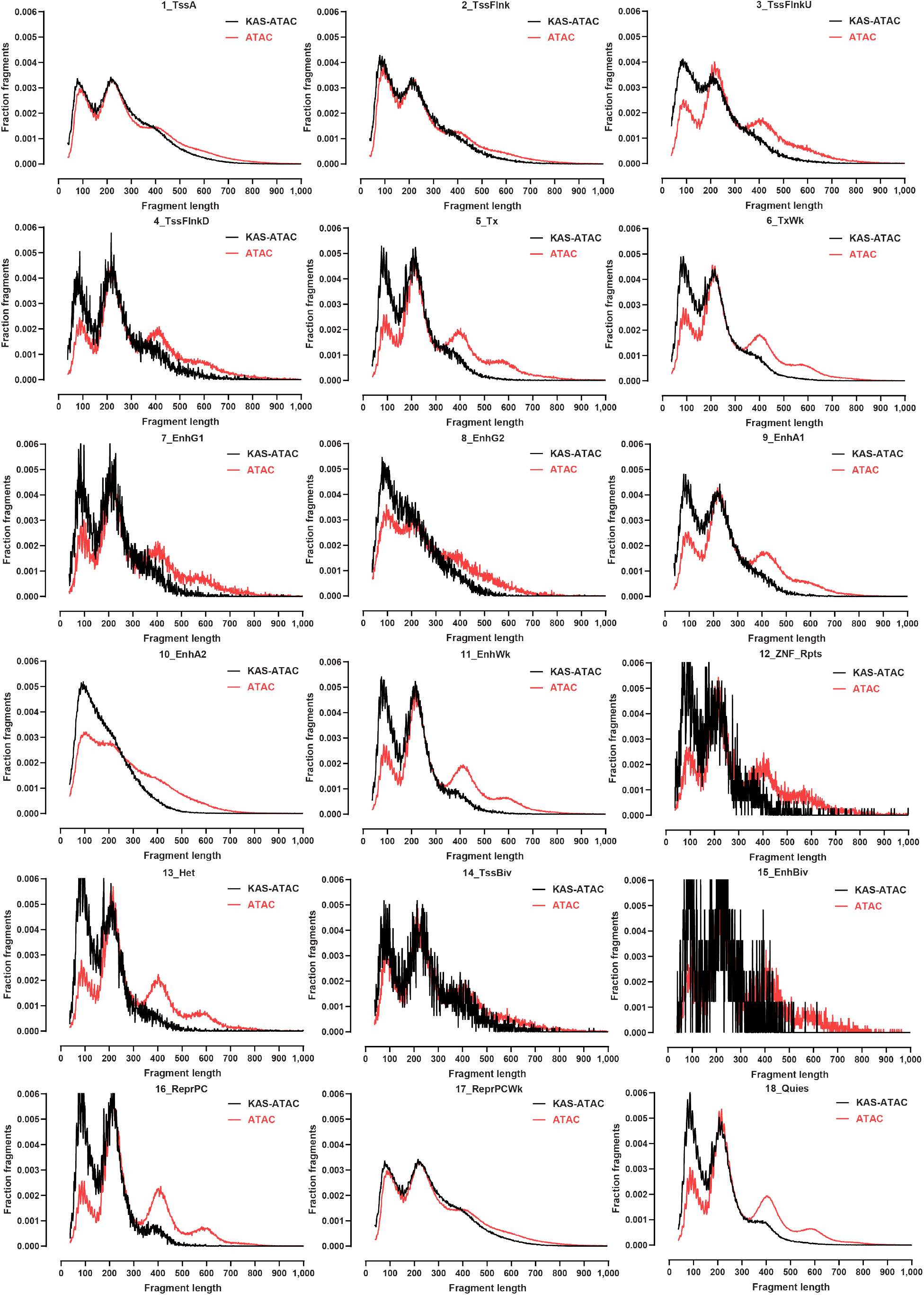
Distribution of KAS-ATAC and ATAC-seq fragment lengths in different chromatin states in HEK293 (replicate 3) cells.

**Supplementary Figure 6:**
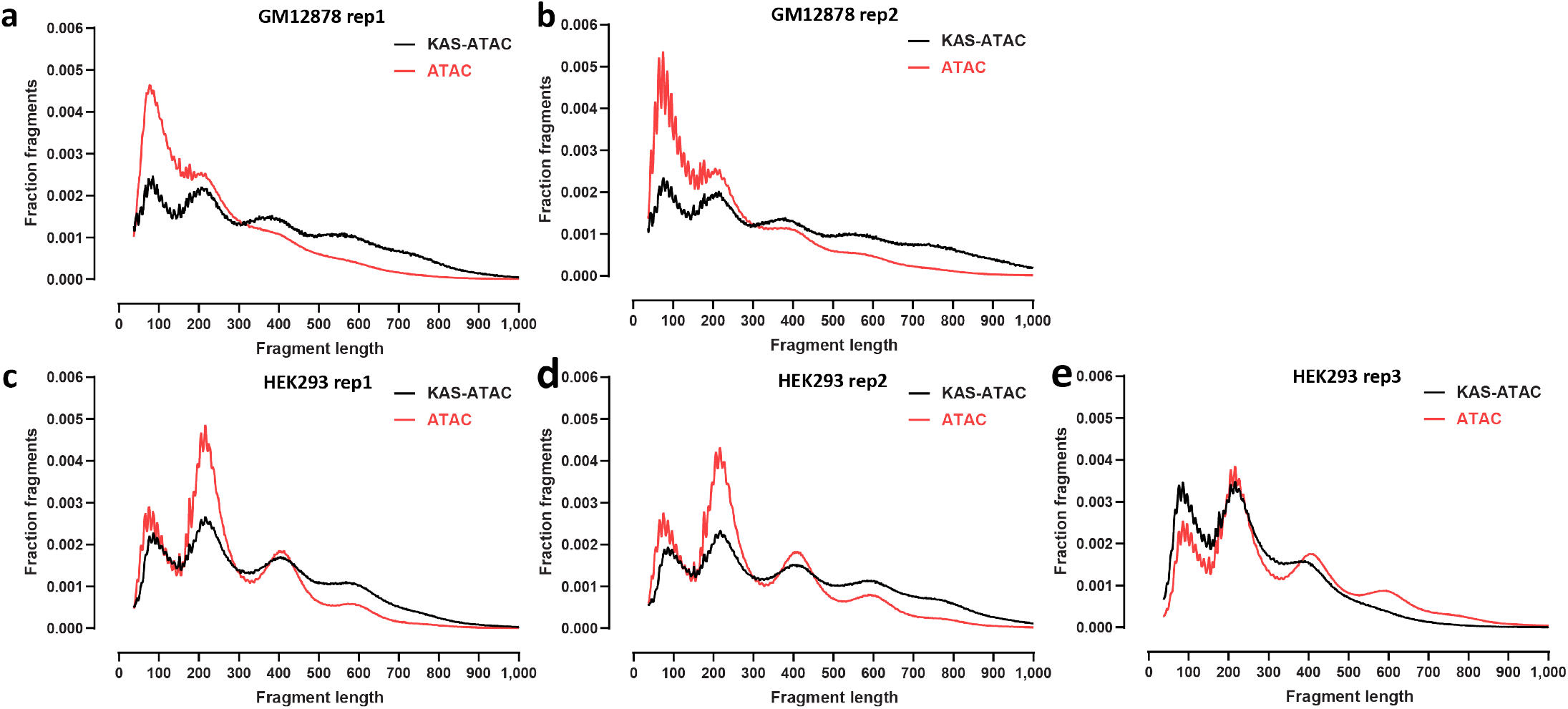
Genome-wide TSS metaprofiles for ATAC-seq and KAS-ATAC libraries. (a) GM12878 cells, replicate 1; (b) GM12878 cells, replicate 2; (c) HEK293 cells, replicate 1; (d) HEK293 cells, replicate 2; (e) HEK293 cells, replicate 3.

**Supplementary Figure 7:**
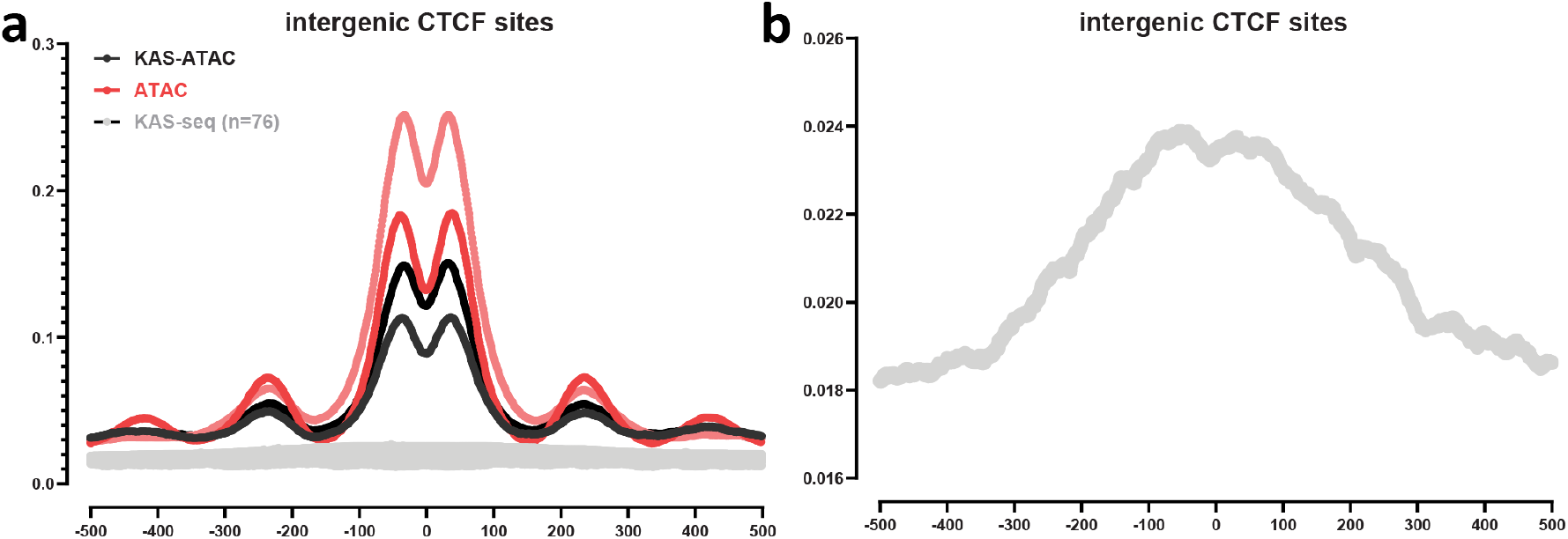
KAS-seq, KAS-ATAC and KAS levels (normalized to RPM) around intergenic (*≥* 1,000 bp away from genes) CTCF sites in HEK293 cells. (a) All three modalities. KAS-seq libraries used are from Marinov & Kim et al. 2023 ^53^ (b) Representative KAS-seq dataset only.

**Supplementary Figure 8:**
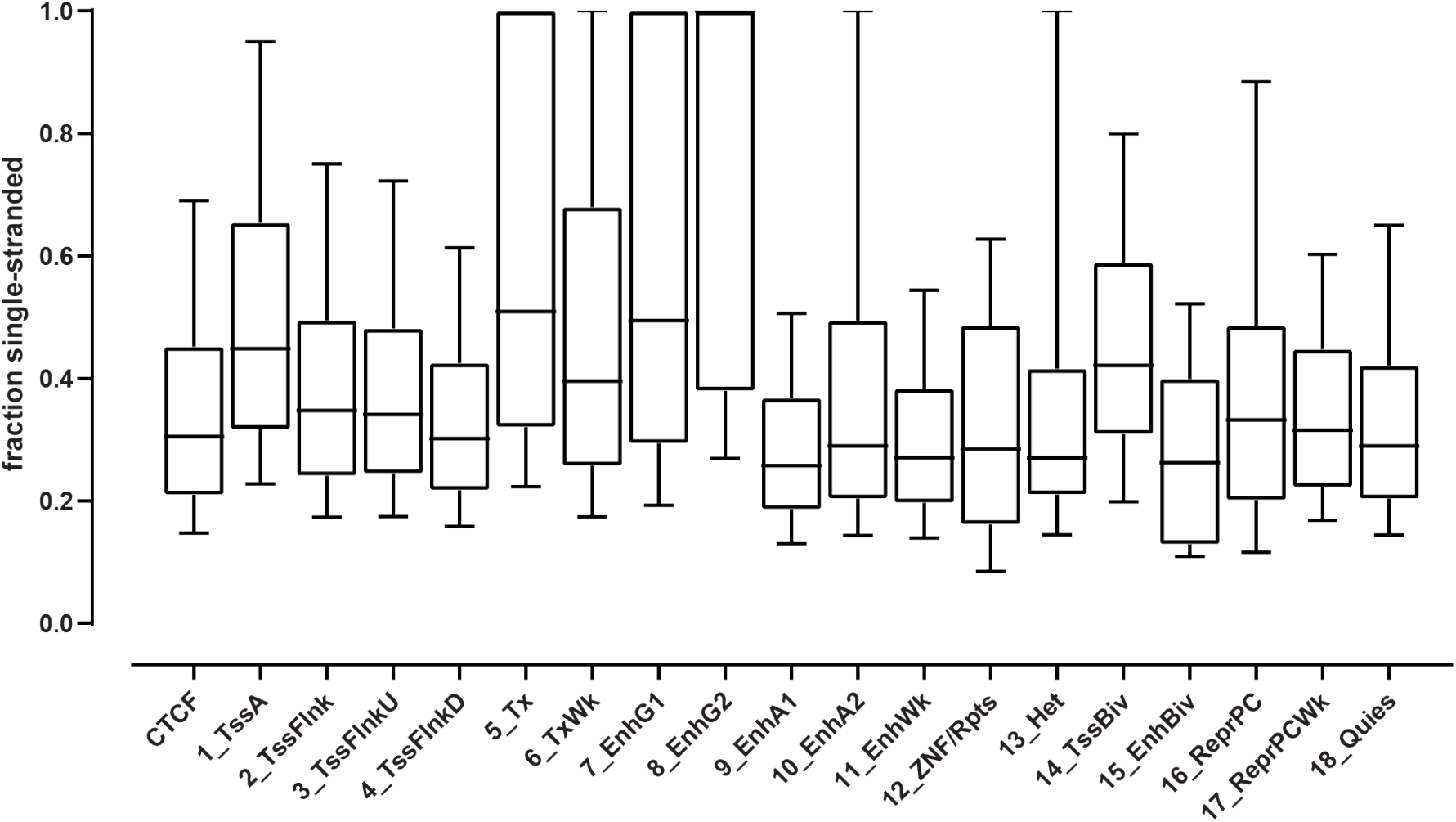
Estimating absolute levels of single-stranded accessible DNA. Distribution of the ratios between the number of KAS-ATAC molecules and ATAC molecules for ATAC-seq peaks found in different chromatin states and for CTCF occupancy sites.

**Supplementary Figure 9:**
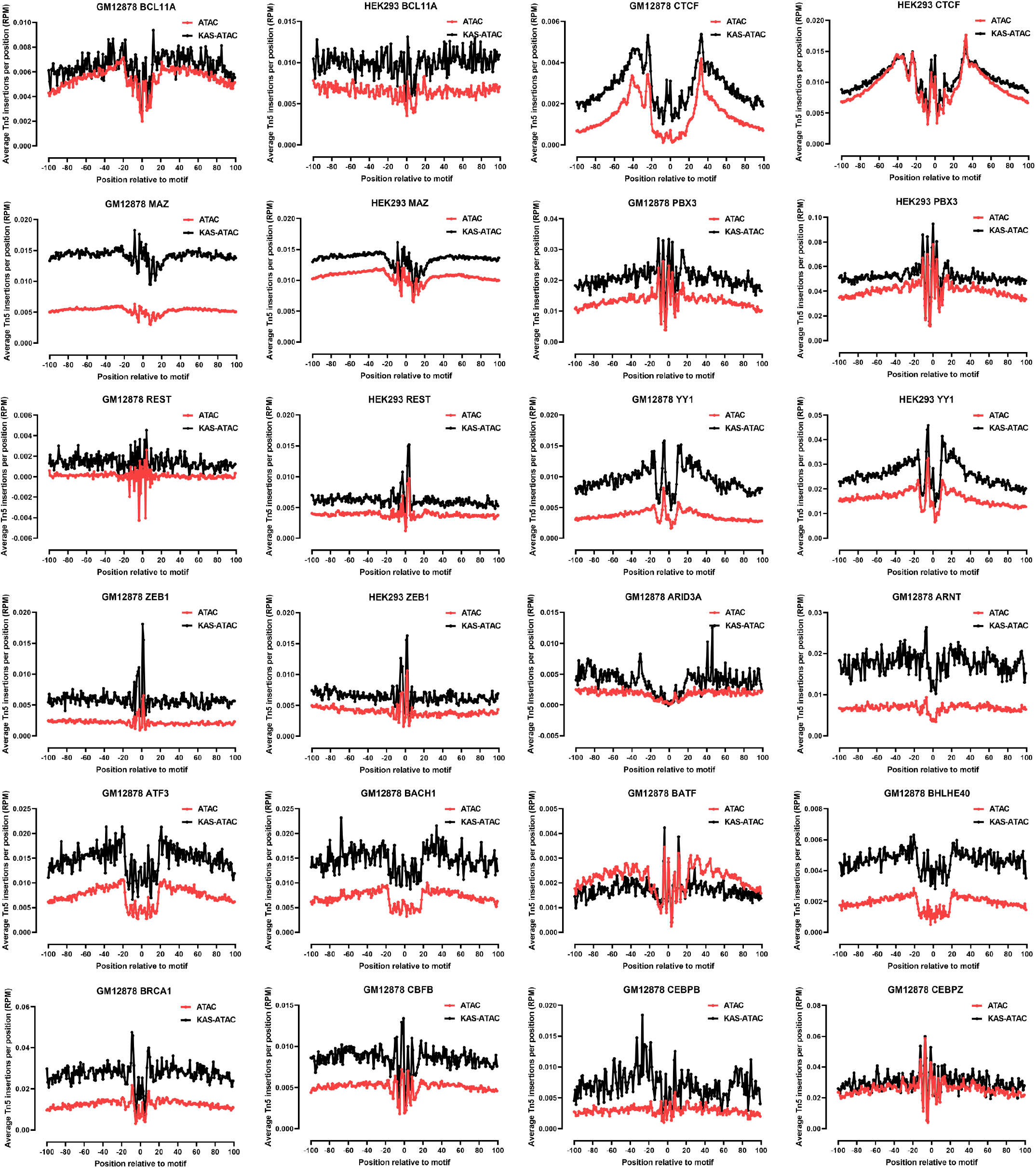
Transcription factor footprints in KAS-ATAC and ATAC datasets. Shown are the average Tn5 insertion profiles (see Methods for details) around the motifs for the indicated factors that are located within ENCODE ChIP-seq peaks for the TF in the given cell line.

**Supplementary Figure 10:**
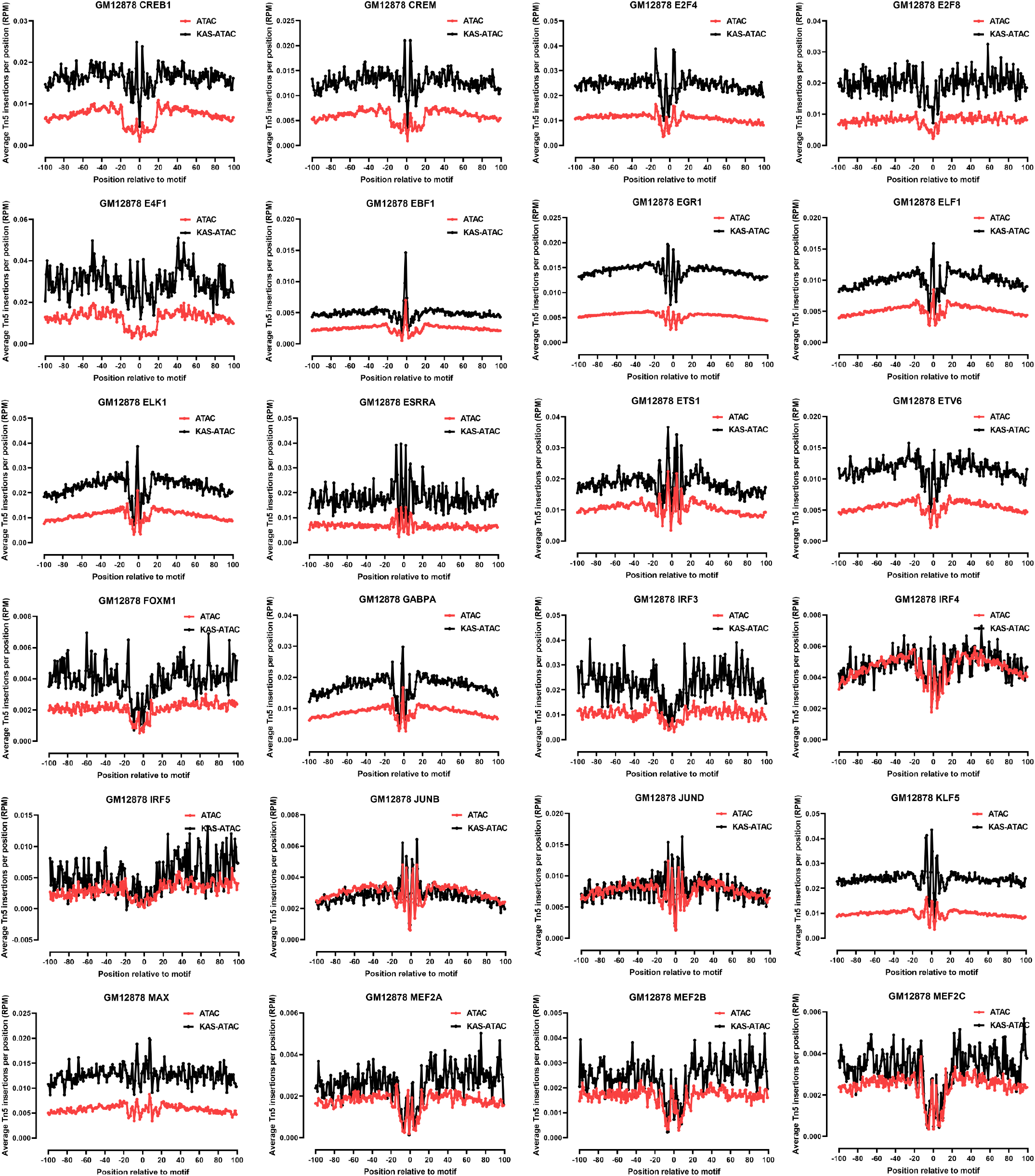
Transcription factor footprints in KAS-ATAC and ATAC datasets. Shown are the average Tn5 insertion profiles (see Methods for details) around the motifs for the indicated factors that are located within ENCODE ChIP-seq peaks for the TF in the given cell line.

**Supplementary Figure 11:**
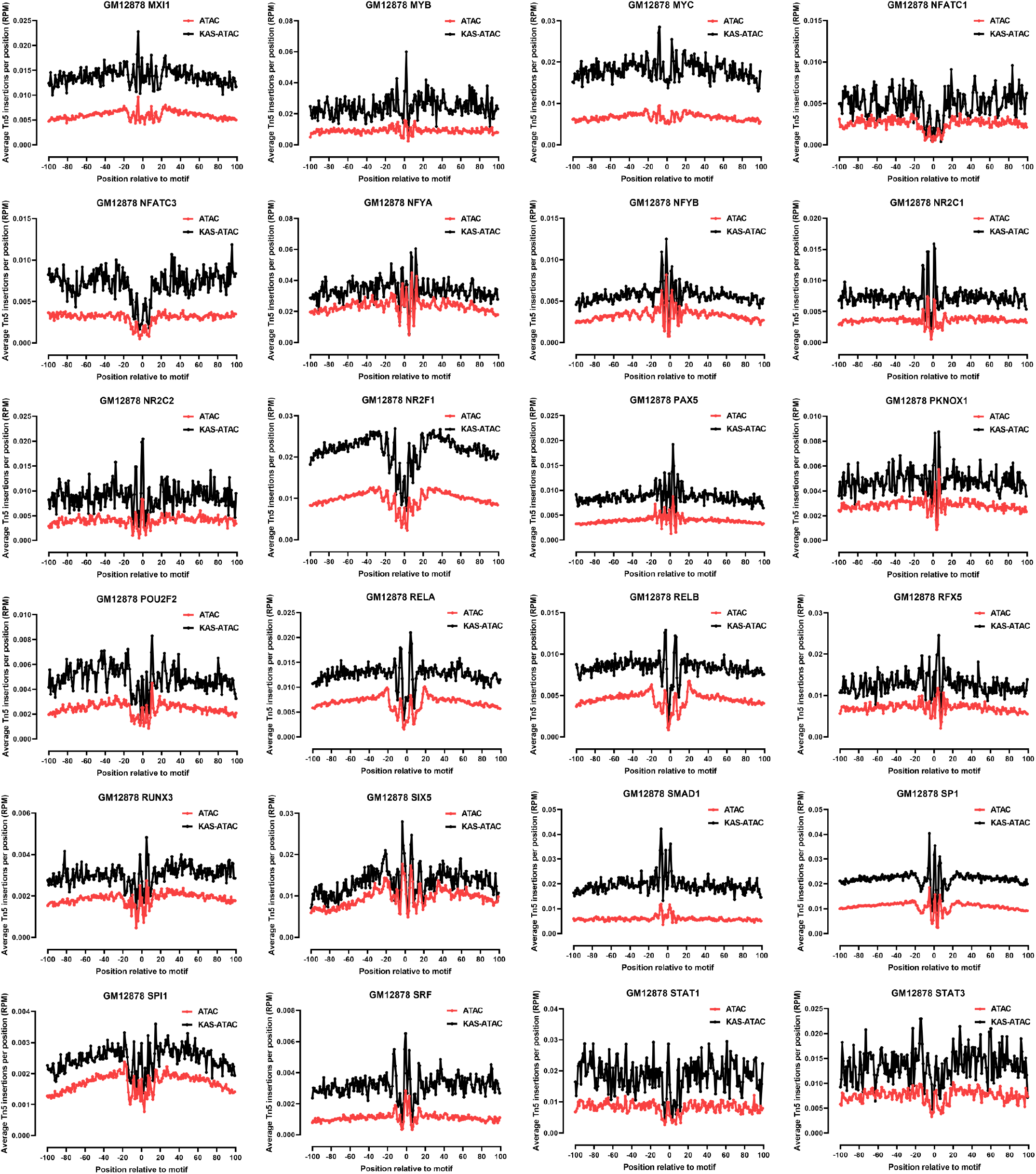
Transcription factor footprints in KAS-ATAC and ATAC datasets. Shown are the average Tn5 insertion profiles (see Methods for details) around the motifs for the indicated factors that are located within ENCODE ChIP-seq peaks for the TF in the given cell line.

**Supplementary Figure 12:**
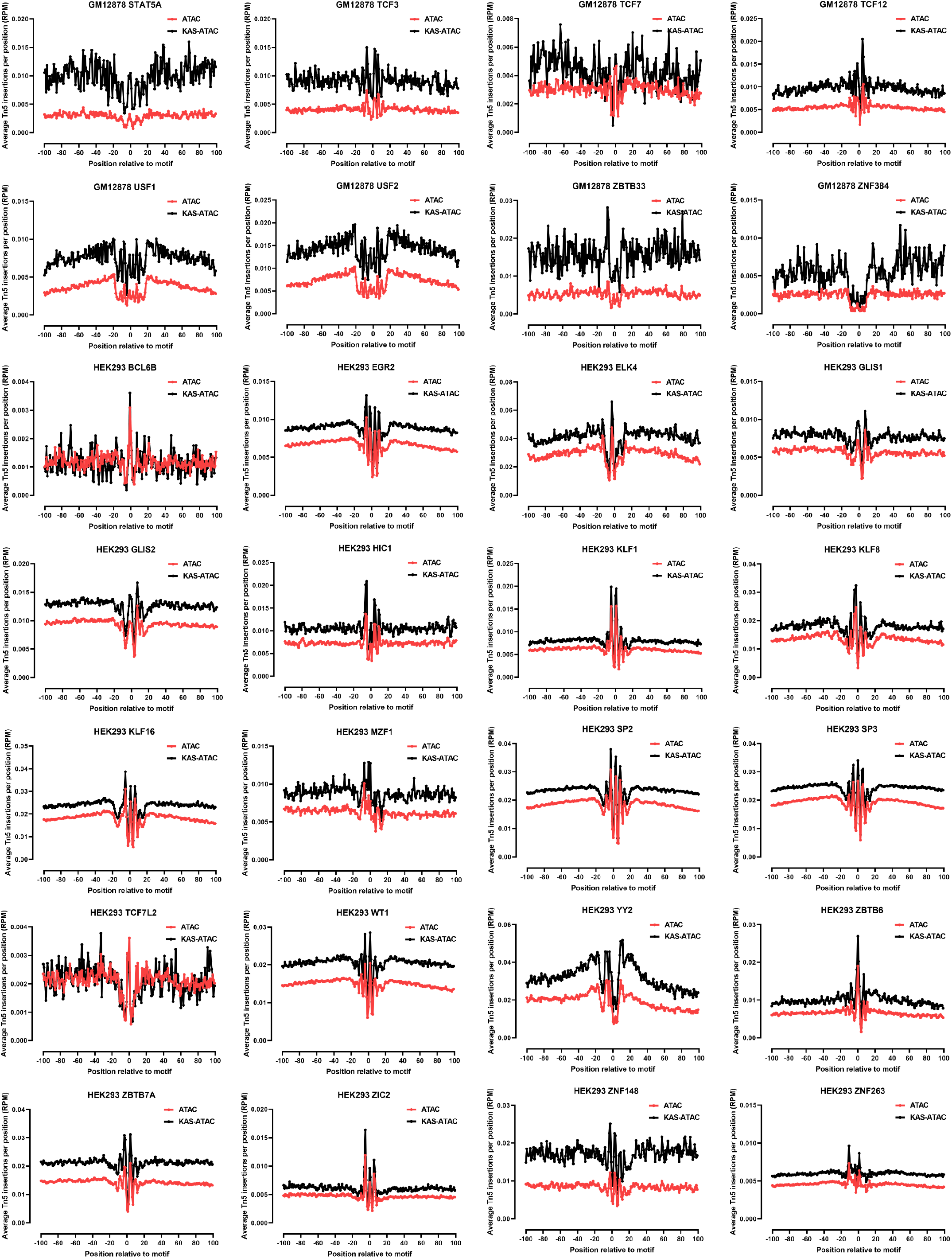
Transcription factor footprints in KAS-ATAC and ATAC datasets. Shown are the average Tn5 insertion profiles (see Methods for details) around the motifs for the indicated factors that are located within ENCODE ChIP-seq peaks for the TF in the given cell line.

## Notes

### Competing Interest Statement

The authors have declared no competing interest.

## References

1. Wu C. 1980. The 5′ ends of Drosophila heat shock genes in chromatin are hypersensitive to DNase I. Nature 286(5776):854–860.

2. Keene MA, Corces V, Lowenhaupt K, Elgin SC. 1981. DNase I hypersensitive sites in Drosophila chromatin occur at the 5′ ends of regions of transcription. Proc Natl Acad Sci U S A 78(1):143–146.

3. McGhee JD, Wood WI, Dolan M, Engel JD, Felsenfeld G. 1981. A 200 base pair region at the 5′ end of the chicken adult β-globin gene is accessible to nuclease digestion. Cell 27(1 Pt 2):45–55.

4. Dorschner MO, Hawrylycz M, Humbert R, Wallace JC, Shafer A, Kawamoto J, Mack J, Hall R, Goldy J, Sabo PJ, Kohli A, Li Q, McArthur M, Stamatoyannopoulos JA. 2004. High-throughput localization of functional elements by quantitative chromatin profiling. Nat Methods 1(3):219–225.

5. Sabo PJ, Kuehn MS, Thurman R, Johnson BE, Johnson EM, Cao H, Yu M, Rosenzweig E, Goldy J, Haydock A, Weaver M, Shafer A, Lee K, Neri F, Humbert R, Singer MA, Richmond TA, Dorschner MO, McArthur M, Hawrylycz M, Green RD, Navas PA, Noble WS, Stamatoyannopoulos JA. 2006. Genome-scale mapping of DNase I sensitivity in vivo using tiling DNA microarrays. Nat Methods 3(7):511–518.

6. Crawford GE, Holt IE, Whittle J, Webb BD, Tai D, Davis S, Margulies EH, Chen Y, Bernat JA, Ginsburg D, Zhou D, Luo S, Vasicek TJ, Daly MJ, Wolfsberg TG, Collins FS. 2006. Genome-wide mapping of DNase hypersensitive sites using massively parallel signature sequencing (MPSS). Genome Res 16(1):123– 131.

7. Boyle AP, Davis S, Shulha HP, Meltzer P, Margulies EH, Weng Z, Furey TS, Crawford GE. 2008. High-resolution mapping and characterization of open chromatin across the genome. Cell 132(2):311–322.

8. Thurman RE, Rynes E, Humbert R, Vierstra J, Maurano MT, Haugen E, Sheffield NC, Stergachis AB, Wang H, Vernot B, Garg K, John S, Sandstrom R, Bates D, Boatman L, Canfield TK, Diegel M, Dunn D, Ebersol AK, Frum T, Giste E, Johnson AK, Johnson EM, Kutyavin T, Lajoie B, Lee BK, Lee K, London D, Lotakis D, Neph S, Neri F, Nguyen ED, Qu H, Reynolds AP, Roach V, Safi A, Sanchez ME, Sanyal A, Shafer A, Simon JM, Song L, Vong S, Weaver M, Yan Y, Zhang Z, Zhang Z, Lenhard B, Tewari M, Dorschner MO, Hansen RS, Navas PA, Stamatoyannopoulos G, Iyer VR, Lieb JD, Sunyaev SR, Akey JM, Sabo PJ, Kaul R, Furey TS, Dekker J, Crawford GE, Stamatoyannopoulos JA. 2012. The accessible chromatin landscape of the human genome. Nature 489(7414):75–82.

9. Kelly TK, Liu Y, Lay FD, Liang G, Berman BP, Jones PA. 2012. Genome-wide mapping of nucleosome positioning and DNA methylation within individual DNA molecules. Genome Res 22(12):2497–2506.

10. Krebs AR, Imanci D, Hoerner L, Gaidatzis D, Burger L, Schübeler D. 2017. Genome-wide Single-Molecule Footprinting Reveals High RNA Polymerase II Turnover at Paused Promoters. Mol Cell 67(3):411–422.e4.

11. Shipony Z, Marinov GK, Swaffer MP, Sinnott-Armstrong NA, Skotheim JM, Kundaje A, Green-leaf WJ. 2020. Long-range single-molecule mapping of chromatin accessibility in eukaryotes. Nat Methods 17(3):319–327.

12. Buenrostro JD, Giresi PG, Zaba LC, Chang HY, Greenleaf WJ. 2013. Transposition of native chromatin for fast and sensitive epigenomic profiling of open chromatin, DNA-binding proteins and nucleosome position. Nat Methods 10(12):1213–1218.

13. Schep AN, Buenrostro JD, Denny SK, Schwartz K, Sherlock G, Greenleaf WJ. 2015. Structured nucleosome fingerprints enable high-resolution mapping of chromatin architecture within regulatory regions. Genome Res 25(11):1757–1770.

14. Hesselberth JR, Chen X, Zhang Z, Sabo PJ, Sandstrom R, Reynolds AP, Thurman RE, Neph S, Kuehn MS, Noble WS, Fields S, Stamatoyannopoulos JA. 2009. Global mapping of protein-DNA interactions in vivo by digital genomic footprinting. Nat Methods 6(4):283–289.

15. Pique-Regi R, Degner JF, Pai AA, Gaffney DJ, Gilad Y, Pritchard JK. 2011. Accurate inference of transcription factor binding from DNA sequence and chromatin accessibility data. Genome Res 21(3):447–455.

16. Neph S, Stergachis AB, Reynolds A, Sandstrom R, Borenstein E, Stamatoyannopoulos JA. 2012. Circuitry and dynamics of human transcription factor regulatory networks. Cell 150(6):1274–1286.

17. Neph S, Vierstra J, Stergachis AB, Reynolds AP, Haugen E, Vernot B, Thurman RE, John S, Sandstrom R, Johnson AK, Maurano MT, Humbert R, Rynes E, Wang H, Vong S, Lee K, Bates D, Diegel M, Roach V, Dunn D, Neri J, Schafer A, Hansen RS, Kutyavin T, Giste E, Weaver M, Canfield T, Sabo P, Zhang M, Balasundaram G, Byron R, MacCoss MJ, Akey JM, Bender MA, Groudine M, Kaul R, Stamatoyannopoulos JA. 2012. An expansive human regulatory lexicon encoded in transcription factor footprints. Nature 489(7414):83–90.

18. Stergachis AB, Neph S, Sandstrom R, Haugen E, Reynolds AP, Zhang M, Byron R, Canfield T, Stelhing-Sun S, Lee K, Thurman RE, Vong S, Bates D, Neri F, Diegel M, Giste E, Dunn D, Vierstra J, Hansen RS, Johnson AK, Sabo PJ, Wilken MS, Reh TA, Treuting PM, Kaul R, Groudine M, Bender MA, Borenstein E, Stamatoyannopoulos JA. 2014. Conservation of trans-acting circuitry during mammalian regulatory evolution. Nature 515(7527):365–370.

19. Vierstra J, Stamatoyannopoulos JA. 2016. Genomic footprinting. Nat Methods 13(3):213–221.

20. Vierstra J, Lazar J, Sandstrom R, Halow J, Lee K, Bates D, Diegel M, Dunn D, Neri F, Haugen E, Rynes E, Reynolds A, Nelson J, Johnson A, Frerker M, Buckley M, Kaul R, Meuleman W, Stamatoyannopoulos JA. 2020. Global reference mapping of human transcription factor footprints. Nature 583(7818):729–736

21. Mortazavi A, Williams BA, McCue K, Schaeffer L, Wold B. 2008. Mapping and quantifying mammalian transcriptomes by RNA-Seq. Nat Methods 5(7):621– 628

22. Nagalakshmi U, Wang Z, Waern K, Shou C, Raha D, Gerstein M, Snyder M. 2008. The transcriptional landscape of the yeast genome defined by RNA sequencing. Science 320(5881):1344–1349.

23. Wilhelm BT, Marguerat S, Watt S, Schubert F, Wood V, Goodhead I, Penkett CJ, Rogers J, Båhler J. 2008. Dynamic repertoire of a eukaryotic transcriptome surveyed at single-nucleotide resolution. Nature 453(7199):1239–1243.

24. Sultan M, Schulz MH, Richard H, Magen A, Klingenhoff A, Scherf M, Seifert M, Borodina T, Soldatov A, Parkhomchuk D, Schmidt D, O’Keeffe S, Haas S, Vingron M, Lehrach H, Yaspo ML. 2008. A global view of gene activity and alternative splicing by deep sequencing of the human transcriptome. Science 321(5891):956–960.

25. Henikoff JG, Belsky JA, Krassovsky K, MacAlpine DM, Henikoff S. 2011. Epigenome characterization at single base-pair resolution. Proc Natl Acad Sci U S A 108:18318–18323

26. Batut P, Dobin A, Plessy C, Carninci P, Gingeras TR. 2013. High-fidelity promoter profiling reveals widespread alternative promoter usage and transposon-driven developmental gene expression. Genome Res 23(1):169–180.

27. Churchman LS, Weissman JS. 2011. Nascent transcript sequencing visualizes transcription at nucleotide resolution. Nature 469(7330):368–373.

28. Core LJ, Waterfall JJ, Lis JT. 2008. Nascent RNA sequencing reveals widespread pausing and divergent initiation at human promoters. Science 322(5909):1845– 1848.

29. Core LJ, Waterfall JJ, Gilchrist DA, Fargo DC, Kwak H, Adelman K, Lis JT. 2012. Defining the status of RNA polymerase at promoters. Cell Rep 2(4):1025– 1035

30. Kwak H, Fuda NJ, Core LJ, Lis JT. 2013. Precise maps of RNA polymerase reveal how promoters direct initiation and pausing. Science 339(6122):950–953.

31. Tome JM, Tippens ND, Lis JT. 2018. Single-molecule nascent RNA sequencing identifies regulatory domain architecture at promoters and enhancers. Nat Genet 50(11):1533–1541

32. Wu T, Lyu R, You Q, He C. 2020. Kethoxal-assisted single-stranded DNA sequencing captures global transcription dynamics and enhancer activity in situ. Nat Methods 17(5):515–523.

33. Bochman ML, Paeschke K, Zakian VA. 2012. DNA secondary structures: stability and function of G-quadruplex structures. Nat Rev Genet 13(11):770– 780.

34. ENCODE Project Consortium, Dunham I, Kundaje A, Aldred SF, Collins PJ, Davis CA, Doyle F, Epstein CB, Frietze S, Harrow J, Kaul R, Khatun J, Lajoie BR, Landt SG, Lee BK, Pauli F, Rosenbloom KR, Sabo P, Safi A, Sanyal A, Shoresh N, Simon JM, Song L, Trinklein ND, Altshuler RC, Birney E, Brown JB, Cheng C, Djebali S, Dong X, Dunham I, Ernst J, Furey TS, Gerstein M, Giardine B, Greven M, Hardison RC, Harris RS, Herrero J, Hoffman MM, Iyer S, Kelllis M, Khatun J, Kheradpour P, Kundaje A, Lassman T, Li Q, Lin X, Marinov GK, Merkel A, Mortazavi A, Parker SC, Reddy TE, Rozowsky J, Schlesinger F, Thurman RE, Wang J, Ward LD, Whitfield TW, Wilder SP, Wu W, Xi HS, Yip KY, Zhuang J, Bernstein BE, Birney E, Dunham I, Green ED, Gunter C, Snyder M, Pazin MJ, Lowdon RF, Dillon LA, Adams LB, Kelly CJ, Zhang J, Wexler JR, Green ED, Good PJ, Feingold EA, Bernstein BE, Birney E, Crawford GE, Dekker J, Elinitski L, Farnham PJ, Gerstein M, Giddings MC, Gingeras TR, Green ED, Guigó R, Hardison RC, Hubbard TJ, Kellis M, Kent WJ, Lieb JD, Margulies EH, Myers RM, Snyder M, Starnatoyannopoulos JA, Tennebaum SA, Weng Z, White KP, Wold B, Khatun J, Yu Y, Wrobel J, Risk BA, Gunawardena HP, Kuiper HC, Maier CW, Xie L, Chen X, Giddings MC, Bernstein BE, Epstein CB, Shoresh N, Ernst J, Kheradpour P, Mikkelsen TS, Gillespie S, Goren A, Ram O, Zhang X, Wang L, Issner R, Coyne MJ, Durham T, Ku M, Truong T, Ward LD, Altshuler RC, Eaton ML, Kellis M, Djebali S, Davis CA, Merkel A, Dobin A, Lassmann T, Mortazavi A, Tanzer A, Lagarde J, Lin W, Schlesinger F, Xue C, Marinov GK, Khatun J, Williams BA, Zaleski C, Rozowsky J, Röder M, Kokocinski F, Abdelhamid RF, Alioto T, Antoshechkin I, Baer MT, Batut P, Bell I, Bell K, Chakrabortty S, Chen X, Chrast J, Curado J, Derrien T, Drenkow J, Dumais E, Dumais J, Duttagupta R, Fastuca M, Fejes-Toth K, Ferreira P, Foissac S, Fullwood MJ, Gao H, Gonzalez D, Gordon A, Gunawardena HP, Howald C, Jha S, Johnson R, Kapranov P, King B, Kingswood C, Li G, Luo OJ, Park E, Preall JB, Presaud K, Ribeca P, Risk BA, Robyr D, Ruan X, Sammeth M, Sandu KS, Schaeffer L, See LH, Shahab A, Skancke J, Suzuki AM, Takahashi H, Tilgner H, Trout D, Walters N, Wang H, Wrobel J, Yu Y, Hayashizaki Y, Harrow J, Gerstein M, Hubbard TJ, Reymond A, Antonarakis SE, Hannon GJ, Giddings MC, Ruan Y, Wold B, Carninci P, Guigó R, Gingeras TR, Rosenbloom KR, Sloan CA, Learned K, Malladi VS, Wong MC, Barber GP, Cline MS, Dreszer TR, Heitner SG, Karolchik D, Kent WJ, Kirkup VM, Meyer LR, Long JC, Maddren M, Raney BJ, Furey TS, Song L, Grasfeder LL, Giresi PG, Lee BK, Battenhouse A, Sheffield NC, Simon JM, Showers KA, Safi A, London D, Bhinge AA, Shestak C, Schaner MR, Kim SK, Zhang ZZ, Mieczkowski PA, Mieczkowska JO, Liu Z, McDaniell RM, Ni Y, Rashid NU, Kim MJ, Adar S, Zhang Z, Wang T, Winter D, Keefe D, Birney E, Iyer VR, Lieb JD, Crawford GE, Li G, Sandhu KS, Zheng M, Wang P, Luo OJ, Shahab A, Fullwood MJ, Ruan X, Ruan Y, Myers RM, Pauli F, Williams BA, Gertz J, Marinov GK, Reddy TE, Vielmetter J, Partridge EC, Trout D, Varley KE, Gasper C, Bansal A, Pepke S, Jain P, Amrhein H, Bowling KM, Anaya M, Cross MK, King B, Muratet MA, Antoshechkin I, Newberry KM, McCue K, Nesmith AS, Fisher-Aylor KI, Pusey B, DeSalvo G, Parker SL, Balasubramanian S, Davis NS, Meadows SK, Eggleston T, Gunter C, Newberry JS, Levy SE, Absher DM, Mortazavi A, Wong WH, Wold B, Blow MJ, Visel A, Pennachio LA, Elnitski L, Margulies EH, Parker SC, Petrykowska HM, Abyzov A, Aken B, Barrell D, Barson G, Berry A, Bignell A, Boychenko V, Bussotti G, Chrast J, Davidson C, Derrien T, Despacio-Reyes G, Diekhans M, Ezkurdia I, Frankish A, Gilbert J, Gonzalez JM, Griffiths E, Harte R, Hendrix DA, Howald C, Hunt T, Jungreis I, Kay M, Khurana E, Kokocinski F, Leng J, Lin MF, Loveland J, Lu Z, Manthravadi D, Mariotti M, Mudge J, Mukherjee G, Notredame C, Pei B, Rodriguez JM, Saunders G, Sboner A, Searle S, Sisu C, Snow C, Steward C, Tanzer A, Tapanan E, Tress ML, van Baren MJ, Walters N, Washieti S, Wilming L, Zadissa A, Zhengdong Z, Brent M, Haussler D, Kellis M, Valencia A, Gerstein M, Raymond A, Guigó R, Harrow J, Hubbard TJ, Landt SG, Frietze S, Abyzov A, Addleman N, Alexander RP, Auerbach RK, Balasubramanian S, Bettinger K, Bhardwaj N, Boyle AP, Cao AR, Cayting P, Charos A, Cheng Y, Cheng C, Eastman C, Euskirchen G, Fleming JD, Grubert F, Habegger L, Hariharan M, Harmanci A, Iyenger S, Jin VX, Karczewski KJ, Kasowski M, Lacroute P, Lam H, Larnarre-Vincent N, Leng J, Lian J, Lindahl-Allen M, Min R, Miotto B, Monahan H, Moqtaderi Z, Mu XJ, O’Geen H, Ouyang Z, Patacsil D, Pei B, Raha D, Ramirez L, Reed B, Rozowsky J, Sboner A, Shi M, Sisu C, Slifer T, Witt H, Wu L, Xu X, Yan KK, Yang X, Yip KY, Zhang Z, Struhl K, Weissman SM, Gerstein M, Farnham PJ, Snyder M, Tenebaum SA, Penalva LO, Doyle F, Karmakar S, Landt SG, Bhanvadia RR, Choudhury A, Domanus M, Ma L, Moran J, Patacsil D, Slifer T, Victorsen A, Yang X, Snyder M, White KP, Auer T, Centarin L, Eichenlaub M, Gruhl F, Heerman S, Hoeckendorf B, Inoue D, Kellner T, Kirchmaier S, Mueller C, Reinhardt R, Schertel L, Schneider S, Sinn R, Wittbrodt B, Wittbrodt J, Weng Z, Whitfield TW, Wang J, Collins PJ, Aldred SF, Trinklein ND, Partridge EC, Myers RM, Dekker J, Jain G, Lajoie BR, Sanyal A, Balasundaram G, Bates DL, Byron R, Canfield TK, Diegel MJ, Dunn D, Ebersol AK, Ebersol AK, Frum T, Garg K, Gist E, Hansen RS, Boatman L, Haugen E, Humbert R, Jain G, Johnson AK, Johnson EM, Kutyavin TM, Lajoie BR, Lee K, Lotakis D, Maurano MT, Neph SJ, Neri FV, Nguyen ED, Qu H, Reynolds AP, Roach V, Rynes E, Sabo P, Sanchez ME, Sandstrom RS, Sanyal A, Shafer AO, Stergachis AB, Thomas S, Thurman RE, Vernot B, Vierstra J, Vong S, Wang H, Weaver MA, Yan Y, Zhang M, Akey JA, Bender M, Dorschner MO, Groudine M, MacCoss MJ, Navas P, Stamatoyannopoulos G, Kaul R, Dekker J, Stamatoyannopoulos JA, Dunham I, Beal K, Brazma A, Flicek P, Herrero J, Johnson N, Keefe D, Lukk M, Luscombe NM, Sobral D, Vaquerizas JM, Wilder SP, Batzoglou S, Sidow A, Hussami N, Kyriazopoulou-Panagiotopoulou S, Libbrecht MW, Schaub MA, Kundaje A, Hardison RC, Miller W, Giardine B, Harris RS, Wu W, Bickel PJ, Banfai B, Boley NP, Brown JB, Huang H, Li Q, Li JJ, Noble WS, Bilmes JA, Buske OJ, Hoffman MM, Sahu AO, Kharchenko PV, Park PJ, Baker D, Taylor J, Weng Z, Iyer S, Dong X, Greven M, Lin X, Wang J, Xi HS, Zhuang J, Gerstein M, Alexander RP, Balasubramanian S, Cheng C, Harmanci A, Lochovsky L, Min R, Mu XJ, Rozowsky J, Yan KK, Yip KY, Birney E. 2012. An integrated encyclopedia of DNA elements in the human genome. Nature 489(7414):57–74.

35. ENCODE Project Consortium, Moore JE, Purcaro MJ, Pratt HE, Epstein CB, Shoresh N, Adrian J, Kawli T, Davis CA, Dobin A, Kaul R, Halow J, Van Nostrand EL, Freese P, Gorkin DU, Shen Y, He Y, Mackiewicz M, Pauli-Behn F, Williams BA, Mortazavi A, Keller CA, Zhang XO, Elhajjajy SI, Huey J, Dickel DE, Snetkova V, Wei X, Wang X, Rivera-Mulia JC, Rozowsky J, Zhang J, Chhetri SB, Zhang J, Victorsen A, White KP, Visel A, Yeo GW, Burge CB, Lécuyer E, Gilbert DM, Dekker J, Rinn J, Mendenhall EM, Ecker JR, Kellis M, Klein RJ, Noble WS, Kundaje A, Guigó R, Farnham PJ, Cherry JM, Myers RM, Ren B, Graveley BR, Gerstein MB, Pennacchio LA, Snyder MP, Bernstein BE, Wold B, Hardison RC, Gingeras TR, Stamatoyannopoulos JA, Weng Z. 2020. Expanded encyclopaedias of DNA elements in the human and mouse genomes. Nature 583(7818):699–710.

36. Marinov GK, Wang YE, Chan D, Wold BJ. 2014. Evidence for site-specific occupancy of the mitochondrial genome by nuclear transcription factors. PLoS One 9(1):e84713.

37. Shadel GS, Clayton DA. 1997. Mitochondrial DNA maintenance in vertebrates. Annu Rev Biochem 66:409–435.

38. Cantatore P, Attardi G. 1980. Mapping of nascent light and heavy strand transcripts on the physical map of HeLa cell mitochondrial DNA. Nucleic Acids Res 8(12):2605–2625.

39. Montoya J, Christianson T, Levens D, Rabinowitz M, Attardi G. 1982. Identification of initiation sites for heavy-strand and light-strand transcription in human mitochondrial DNA. Proc Natl Acad Sci 79(23):7195– 7199.

40. Ernst J, Kellis M. 2012. ChromHMM: automating chromatin-state discovery and characterization. Nat Methods 9(3):215–216.

41. Love MI, Huber W, Anders S. 2014. Moderated estimation of fold change and dispersion for RNA-seq data with DESeq2. Genome Biol 15(12):550.

42. Bell AC, West AG, Felsenfeld G. 1999. The protein CTCF is required for the enhancer blocking activity of vertebrate insulators. Cell 98(3):387–399.

43. Fu Y, Sinha M, Peterson CL, Weng Z. 2008. The insulator binding protein CTCF positions 20 nucleosomes around its binding sites across the human genome. PLoS Genet 4(7):e1000138

44. Wulfridge P, Yan Q, Rell N, Doherty J, Jacobson S, Offley S, Deliard S, Feng K, Phillips-Cremins JE, Gardini A, Sarma K. 2023. G-quadruplexes associated with R-loops promote CTCF binding. Mol Cell 83(17):3064–3079.e5

45. Seila AC, Calabrese JM, Levine SS, Yeo GW, Rahl PB, Flynn RA, Young RA, Sharp PA. 2008. Divergent transcription from active promoters. Science 322(5909):1849–1851.

46. Daley T, Smith AD. 2013. Predicting the molecular complexity of sequencing libraries. Nat Methods 10(4):325–327.

47. Weirauch MT, Yang A, Albu M, Cote AG, Montenegro-Montero A, Drewe P, Najafabadi HS, Lambert SA, Mann I, Cook K, Zheng H, Goity A, van Bakel H, Lozano JC, Galli M, Lewsey MG, Huang E, Mukherjee T, Chen X, Reece-Hoyes JS, Govindarajan S, Shaulsky G, Walhout AJ, Bouget FY, Ratsch G, Larrondo LF, Ecker JR, Hughes TR. 2014. Determination and inference of eukaryotic transcription factor sequence specificity. Cell 158(6):1431–1443.

48. Sung MH, Baek S, Hager GL. 2016. Genome-wide footprinting: ready for prime time? Nat Methods 13(3):222–228.

49. Zhang H, Lu T, Liu S, Yang J, Sun G, Cheng T, Xu J, Chen F, Yen K. 2021. Comprehensive understanding of Tn5 insertion preference improves transcription regulatory element identification. NAR Genom Bioinform 3(4):lqab094

50. Li Z, Schulz MH, Look T, Begemann M, Zenke M, Costa IG. 2019. Identification of transcription factor binding sites using ATAC-seq. Genome Biol 20(1):45.

51. Baek S, Goldstein I, Hager GL. 2017. Bivariate Genomic Footprinting Detects Changes in Transcription Factor Activity. Cell Rep 19(8):1710–1722.

52. Bentsen M, Goymann P, Schultheis H, Klee K, Petrova A, Wiegandt R, Fust A, Preussner J, Kuenne C, Braun T, Kim J, Looso M. 2020. ATAC-seq footprinting unravels kinetics of transcription factor binding during zygotic genome activation. Nat Commun 11(1):4267

53. Marinov GK, Kim SH, Bagdatli ST, Higashino SI, Trevino AE, Tycko J, Wu T, Bintu L, Bassik MC, He C, Kundaje A, Greenleaf WJ. 2023. CasKAS: direct profiling of genome-wide dCas9 and Cas9 specificity using ssDNA mapping. Genome Biol 24(1):85

54. Picelli S, Björklund AK, Reinius B, Sagasser S, Winberg G, Sandberg R. 2014. Tn5 transposase and tagmentation procedures for massively scaled sequencing projects. Genome Res 24(12):2033–2040.

55. Corces MR, Trevino AE, Hamilton EG, Greenside PG, Sinnott-Armstrong NA, Vesuna S, Satpathy AT, Rubin AJ, Montine KS, Wu B, Kathiria A, Cho SW, Mumbach MR, Carter AC, Kasowski M, Orloff LA, Risca VI, Kundaje A, Khavari PA, Montine TJ, Greenleaf WJ, Chang HY. 2017. An improved ATAC-seq protocol reduces background and enables interrogation of frozen tissues. Nat Methods 14(10):959–962.

56. Marinov GK, Shipony Z, Kundaje A, Greenleaf WJ. 2023. Genome-Wide Mapping of Active Regulatory Elements Using ATAC-seq. Methods Mol Biol 2611:3– 19.

57. Marinov GK, Shipony Z. 2021. Interrogating the Accessible Chromatin Landscape of Eukaryote Genomes Using ATAC-seq. Methods Mol Biol 2243:183–226.

58. Langmead B, Trapnell C, Pop M, Salzberg SL. 2009. Ultrafast and memory-efficient alignment of short DNA sequences to the human genome. Genome Biol 10(3):R25.

59. Feng J, Liu T, Qin B, Zhang Y, Liu XS. 2012. Identifying ChIP-seq enrichment using MACS. Nat Protoc 7(9):1728–1740.

60. Grant CE, Bailey TL, Noble WS. 2011. FIMO: scanning for occurrences of a given motif. Bioinformatics 27(7):1017–1018.

